# Cultured pluripotent planarian stem cells retain potency and express proteins from exogenously introduced mRNAs

**DOI:** 10.1101/573725

**Authors:** Kai Lei, Sean A. McKinney, Eric J. Ross, Heng-Chi Lee, Alejandro Sánchez Alvarado

## Abstract

Planarians possess naturally occurring pluripotent adult somatic stem cells (neoblasts) required for homeostasis and whole–body regeneration. However, methods for culturing neoblasts are currently unavailable, hindering both mechanistic studies of potency and the development of transgenic tools. We report the first robust methodologies for culturing and delivering exogenous mRNA into neoblasts. We identified culture media for maintaining neoblasts in vitro, and showed via transplantation that the cultured stem cells retained pluripotency. By modifying standard flow cytometry methods, we developed a new procedure that significantly improved yield and purity of neoblasts. These methods facilitated the successful introduction and expression of exogenous mRNAs in neoblasts, overcoming a key hurdle impeding the application of transgenics in planarians. The tissue culture advances reported here create new opportunities to advance detailed mechanistic studies of adult stem cell pluripotency in planarians, and provide a systematic methodological framework to develop cell culture techniques for other emerging research organisms.

## Introduction

Stem cell pluripotency remains an important and still unresolved problem in biology. Several systems have been established to study pluripotency regulation in germlines, embryonic, and induced pluripotent stem cells ^1–4^. However, no naturally occurring adult pluripotent stem cells have been identified in traditional model systems, including round worms, flies, fishes, and mice. Unlike traditional research organisms, planarians harbor an abundant population of adult stem cells collectively known as neoblasts. These cells are characteristic of flatworms and acoels ^5^, and in planarians include a subpopulation of pluripotent stem cells termed clonogenic neoblasts ^6–8^. Neoblasts confer planarians with remarkable regenerative abilities and a seemingly limitless capacity for tissue homeostasis. Of the many freshwater planarian species known to exist, *Schmidtea mediterranea* has become one of the most widely studied ^9^. Planarians thus provide a unique context in which to explore how nature has solved the complex problem of maintaining stem cell pluripotency in a long-lived adult animal.

Expression of conserved genes regulating pluripotency have been identified in planarian neoblasts and functionally studied using RNA interference ^10–13^. However, due in part to the lack of methodologies for cell culture, exogenous gene expression, and transgenesis in planarians, the mechanisms regulating the pluripotency of these adult stem cells *in vivo* are poorly understood. Therefore, developing planarian transgenesis is of great significance ^14^. A review of the history of cell culture methodologies and attempts to develop transgenics in planarians indicated that successful neoblast culture may be a critical first step to develop transgenic methodologies in planarians.

Transgenic approaches typically take advantage of either early stage embryos or cultured stem cells. Invertebrates, such as *Caenorhabditis elegans*, *Drosophila melanogaster*, Hydra, *Nematostella vectensis*, and the flatworm *Macrostomum lignano*, have large syncytial germ cells or embryos, respectively, that are highly amenable to genetic manipulation ^14–18^. In vertebrates, such as mice, both zygotes and cultured embryonic stem cells are used to deliver exogenous genetic material^19^. Unlike these research organisms, planarians do not possess large, easily accessible germ cells or early-stage blastomeres amenable to manipulation or transplantation. Instead, in asexually reproducing planarians, neoblasts are the only known proliferating cells in the animal ^20^. Neoblasts from one animal can be readily transplanted into a host devoid of its own endogenous neoblasts after lethal irradiation, resulting in neoblast repopulation and host rescue within 1 month ^6, 8^. Thus, introduction of exogenous DNA into cultured neoblasts prior to transplantation is a potential strategy to produce transgenic planarians. Cultured neoblasts would also be ideal for rapidly screening conditions for delivering and expressing exogenous mRNA or DNA. Therefore, we aimed to establish a robust method for culturing pluripotent neoblasts that may allow rapid screening of conditions for delivering and expressing exogenous mRNA or DNA.

Previous efforts to culture planarian cells were conducted at a time when our mechanistic understanding of neoblast self-renewal and heterogeneity were limited ^21–24^. In some of these studies, cells with gross morphology similar to neoblasts survived in an isotonic medium for a couple of weeks, yet neither functional nor molecular tests on the cultured cells were performed, leaving an open question as to their identity and potency ^23^. Since then, the pan-neoblast marker *smedwi-1* (a homolog of the Argonaute family of proteins) was identified ^25^, allowing us to molecularly define and visualize neoblasts using gene expression profiling or whole mount in situ hybridization (ISH). Techniques that enrich neoblasts using flow sorting have also been developed. A cell cycle-based flow sorting method using Hoechst 33342 staining has been used to isolate S and G2/M cell cycle phase neoblasts (X1 cells; nearly 90% of X1 cells are *smedwi-1+* neoblasts) ^25, 26^. However, Hoechst 33342 is cytotoxic, and X1 neoblasts cannot proliferate *in vivo* after transplantation into lethally irradiated planarians lacking stem cells. To solve this technical limitation, a DNA dye free back-gating strategy using forward scatter (size) and side scatter (complexity) was shown to enrich for a heterogeneous cell population containing neoblasts (X1(FS)) ^8^. Unlike X1 neoblasts, X1(FS) neoblasts proliferate and successfully rescue lethally irradiated planarians upon transplantation making the X1(FS) population suitable for the development of an *in vitro* neoblast culture protocol ^8^. When considered alongside the formulation of new types of cell culture media ^23, 27^, these advances provide a groundwork to attempt establishing new, robust methods for *in vitro* culture of pluripotent and transplantation-competent neoblasts.

In this study, we performed an unbiased screen of 23 different formulations of cell culture media to identify the best nutrient conditions for flow cytometrically isolated neoblasts. Cell morphology, viability, percentage of *smedwi-1*+ cells, clonogenic capacity after transplantation, and rescue efficiency were assayed to identify the optimal conditions for culturing pluripotent neoblasts. Importantly, time-lapse imaging captured neoblast division for the first time in culture in real-time. Moreover, a novel neoblast isolation method using the vital dye SiR-DNA was developed, improving the purification yields for neoblasts relative to X1(FS), while preserving the clonogenic and rescue capacity of neoblasts following transplantation. Finally, we developed electroporation conditions that can deliver exogenous mRNA into cultured neoblasts providing unambiguous evidence that exogenous mRNAs can be expressed, albeit with low efficiency, in cultured neoblasts. Cumulatively, our work provides a foundation for developing long-term neoblast culture methods and, ultimately, transgenic planarians. It also provides a systematic methodological framework that may be applied to the development of cell culture techniques in other invertebrate research organisms.

## Results

### Identification of seven culture conditions that maintain viable dividing neoblasts

To test different culture conditions for neoblasts, we first used an established back-gating method to sort X1(FS) cells, which contain approximately 23.4%±2.5% neoblasts (*smedwi-1+*) (Fig. 1a-c). We then systematically screened 23 different types of media, representing most commercially available or previously reported formulations in the prescence (+) or absence (-) of 5% CO_2_ (Supplementary Table 1) ^23, 24^. To assess each culture condition, five criteria were assayed: 1) cell morphology and viability (viability); 2) percentage of *smedwi-1*+ cells maintained in culture (%neoblasts); 3) cell division; 4) clonogenic capacity after transplantation (colony expansion); and 5) rescue efficiency in lethally irradiated planarians (pluripotency) (Fig. 1a).

**Figure 1.**
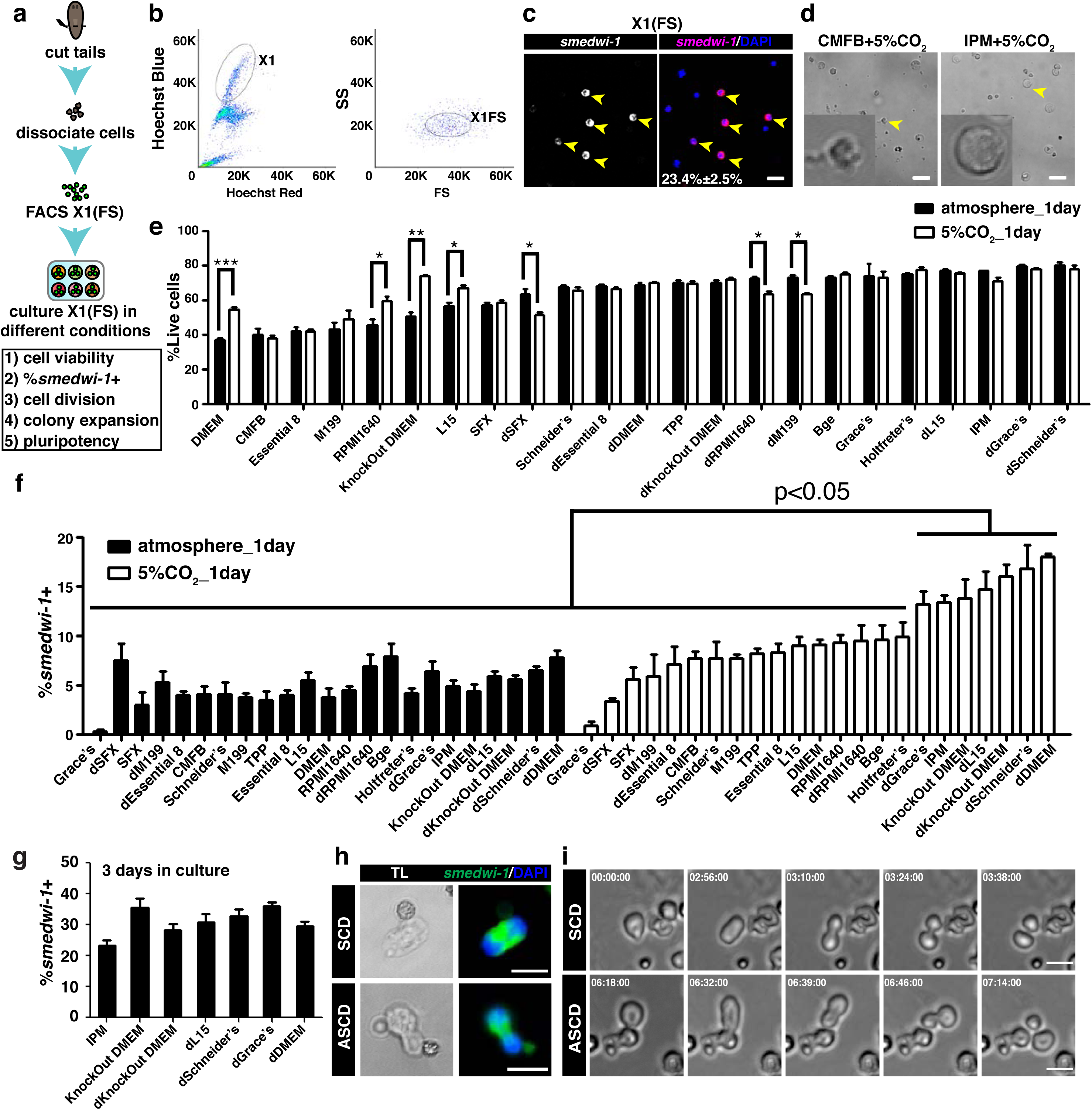
Systematic screen identifies cell culture conditions for maintaining X1(FS) neoblasts *in vitro*. **(a)** Flowchart illustrating steps of X1(FS) cell culture and criteria used to identify best culture condition for neoblasts: cell viability, percentage of *smedwi-1*+ neoblasts (%*smedwi-1*+), cell division *in vitro*, colony expansion after transplantation, and rescue efficiency of irradiated hosts after transplantation (pluripotency). **(b)** Plots showing the FACS gating to sort X1(FS) cells. **(c)** Representative images showing *smedwi-1+* neoblasts among the sorted X1(FS) cells. Scale bar, 20 µm. X1(FS) cells consistently contains 23.4%±2.5% neoblasts in total DAPI+ cells scored. Three replicates were assayed, n=100 to 150. **(d)** Representative images of cell morphologies observed after 1 day of culture +5% CO_2_, including poor cell morphology in CMFB and healthy cell morphology in IPM (arrowheads). Scale bar, 20 µm. **(e)** Percentages of live cells (Propidium Iodide-negative) among 23 media, +/− 5% CO_2_, after 1 day of culture. Knockout DMEM + 5% CO_2_ yielded best overall cell viability. Three replicates were assayed, n=500 to 1200. **(f)** Percentage of *smedwi-1+* neoblasts after 1 day of culture under indicated conditions. Significantly more *smedwi-1*+ neoblasts were maintained in seven media + 5% CO_2_ compared to all other conditions. Three replicates were assayed, n>500. **(g)** Percentage of *smedwi-1+* neoblasts after 3 days of culture in indicated media + 5% CO_2_. **(h)** Representative images of dividing cells undergoing either symmetric cell division (SCD) or asymmetric cell division (ASCD). Scale bar, 10 µm. **(i)** Time-lapse images of dividing cells undergoing either SCD or ASCD in IPM + 5% CO_2_. Scale bar, 10 µm. Both SCD and ASCD can be observed in ∼300 X1(FS) cells cultured in IPM, KnockOut DMEM, and dL15 + 5% CO_2_.

After 1 day of culture, cell morphologies were observed using transmitted light microscopy. Cells cultured in CMFB +/− 5% CO_2_ displayed abnormally roughened cell morphologies accompanied by abundant cellular debris in the plate, suggesting poor viability (Fig. 1d). In contrast, cells in all other conditions, such as IPM +/− 5% CO_2_, had normal morphology, suggesting high viability (Fig. 1d). Cells in Leibovitz’s L-15 medium (L15) without 5% CO_2_ extended long processes that were visible even after 6 days of culture (Supplementary Fig. 1), suggesting neuronal differentiation as previously observed in cultured *Caenorhabditis elegans* embryonic cells ^28^.

To measure viability, cells cultured for 1 day were stained with propidium iodide (PI), which labels the DNA of dying cells, and the percentage of PI negative cells was determined using flow cytometry. Consistent with the microscopic evaluation, cells cultured in CMFB displayed poor survival +/− 5% CO_2_ (>60% dead cells) (Fig. 1e). In fact, of all media conditions tested, seventeen yielded a viability of at least 60% (Fig. 1e), with only seven of the media performing significantly different in the presence and absence of 5% CO_2_ (Fig. 1e).

To determine what proportion of viable cells were neoblasts after 24 hours of culture, we quantified the number of *smedwi-1*+ X1(FS) cells by fluorescent *in situ* hybridization (FISH). Importantly, all cultures without 5% CO_2_ maintained fewer *smedwi-1*+ neoblasts compared to those cultured in the presence of 5% CO_2_, except for diluted (d) SFX (Fig. 1f). Furthermore, of the 5% CO_2_ cultures, seven media maintained significantly more *smedwi-1*+ neoblasts than all other culture conditions, including dGrace’s medium, IPM, KnockOut DMEM, dL15 medium, dKnockOut DMEM, dSchneider’s medium, and dDMEM (Fig. 1f). Because dSFX without 5% CO_2_ failed to support neoblast culture as well as the seven media with 5% CO_2_ we identified above (Fig. 1e), we did not explore it further in this study. This result was supported by co-staining cells cultured in IPM + 5% CO_2_ with *smedwi-1* and the apoptotic/dead-cell marker, Annexin V (Supplementary Fig. 2); no co-labeling was observed, indicating that neoblasts were viable (Supplementary Fig. 2). Consistently, cell viability in these seven media + 5% CO_2_ was consistently greater than 60% (Fig. 1e). We next examined whether *smedwi-1+* neoblasts were maintained after 3 days in culture using these seven media + 5% CO_2_, and observed persistent *smedwi-1+* cells in all culture conditions tested (Fig. 1g). Thus, neoblasts can be maintained for at least 3 days *in vitro*. Therefore, we focused on testing dGrace’s, IPM, KnockOut DMEM, dL15, dKnockOut DMEM, dSchneider’s, and dDMEM media in subsequent optimization experiments.

Next, we assessed whether cultured neoblasts were capable of dividing *in vitro.* Although an obvious increase in cell number was not noticed, low levels of both symmetric and asymmetric neoblast divisions were observed in 1 day cultured cells, as judged by cell pair size and distribution of *smedwi-1* transcripts (Fig. 1h)^13^. Confirmation that neoblasts can divide *in vitro* was obtained using time-lapse imaging microscopy to record the behavior of X1(FS) cells in culture. Both symmetric and asymmetric cell divisions were observed within the first 24 hours after culture (Fig. 1i and Supplementary Movies 1 and 2) in IPM, KnockOut DMEM, and dL15 medium, but not in the other four media tested (Fig. 1i). Consistently, the percentage of *PCNA*+ cells in the cultures of IPM, KnockOut DMEM, and dL15 medium were significantly higher than those in CMFB, Schneider’s, and DMEM medium (Supplementary Fig. 3). Even though we cannot exclude the possibility that these conditions only allow neoblasts in M phase to complete the cell cycle, to our knowledge, this is the first time that neoblast divisions have been observed and recorded *in vitro*. These results suggest that a fraction of X1(FS), *smedwi-1*+ cells can execute cell division within 24 hours after isolation in culture.

### Cultured neoblasts maintain clonogenic capacity

To determine if X1(FS) neoblasts could divide *in vivo* following *in vitro* culture, we next examined their clonogenic capacity following transplantation into lethally irradiated planarians, an experimental manipulation that normally leads to robust neoblast expansion (Fig. 2a). Serial cell dilution experiments indicated that 1,000 freshly collected X1(FS) cells undergo consistent colony expansion in ≥ 80% hosts upon bulk cell transplantation (Supplementary Fig. 4). Considering the rate of cell death in culture, 5,000 X1(FS) cells were cultured for each test condition to ensure that enough cells were viable at the time of transplant. We transplanted X1(FS) cells cultured in the seven different media + 5% CO_2_ that showed higher than 15% *smedwi-1*+ cells (Figure 1f) for 1, 2, or 3 days. At 8 days post-transplantation (dpt), the presence or absence of *smedwi-1*+ neoblast colonies and the number of *smedwi-1*+ neoblasts in each colony were determined. All X1(FS) neoblasts cultured for 1 or 2 days efficiently proliferated *in vivo*, except for those cultured in dGrace’s medium + 5% CO_2_ (Fig. 2b–d). By comparing the number of *smedwi-1*+ neoblasts in each transplant, X1(FS) cells cultured for 1 day in either IPM or KnockOut DMEM formed the largest colonies *in vivo* (Fig. 2b, d). X1(FS) cells cultured for 2 days displayed decreased expansion potential in all conditions, but all were still capable of forming colonies *in vivo* with the exception of those cultured in dGrace’s medium + 5% CO_2_. X1(FS) cells cultured for 3 days were largely incapable of forming colonies following transplantation, though small colonies formed from cells cultured in dSchneider and dL15 media (Fig. 2c, d). In summary, IPM and KnockOut DMEM performed best following the first day in culture, but performed similarly to dKnockOut DMEM, dSchneider’s, dL15, and dDMEM after two days of culture. In addition, we observed that clonogenic capacity of X1(FS) neoblasts diminished greatly following three days in culture, regardless of the media used. These results suggest that IPM, KnockOut DMEM, dL15, dKnockOut DMEM, dSchneider’s, and dDMEM are capable of maintaining the proliferation potential of neoblasts for up to two days in culture in the presence of 5% CO_2_.

**Figure 2.**
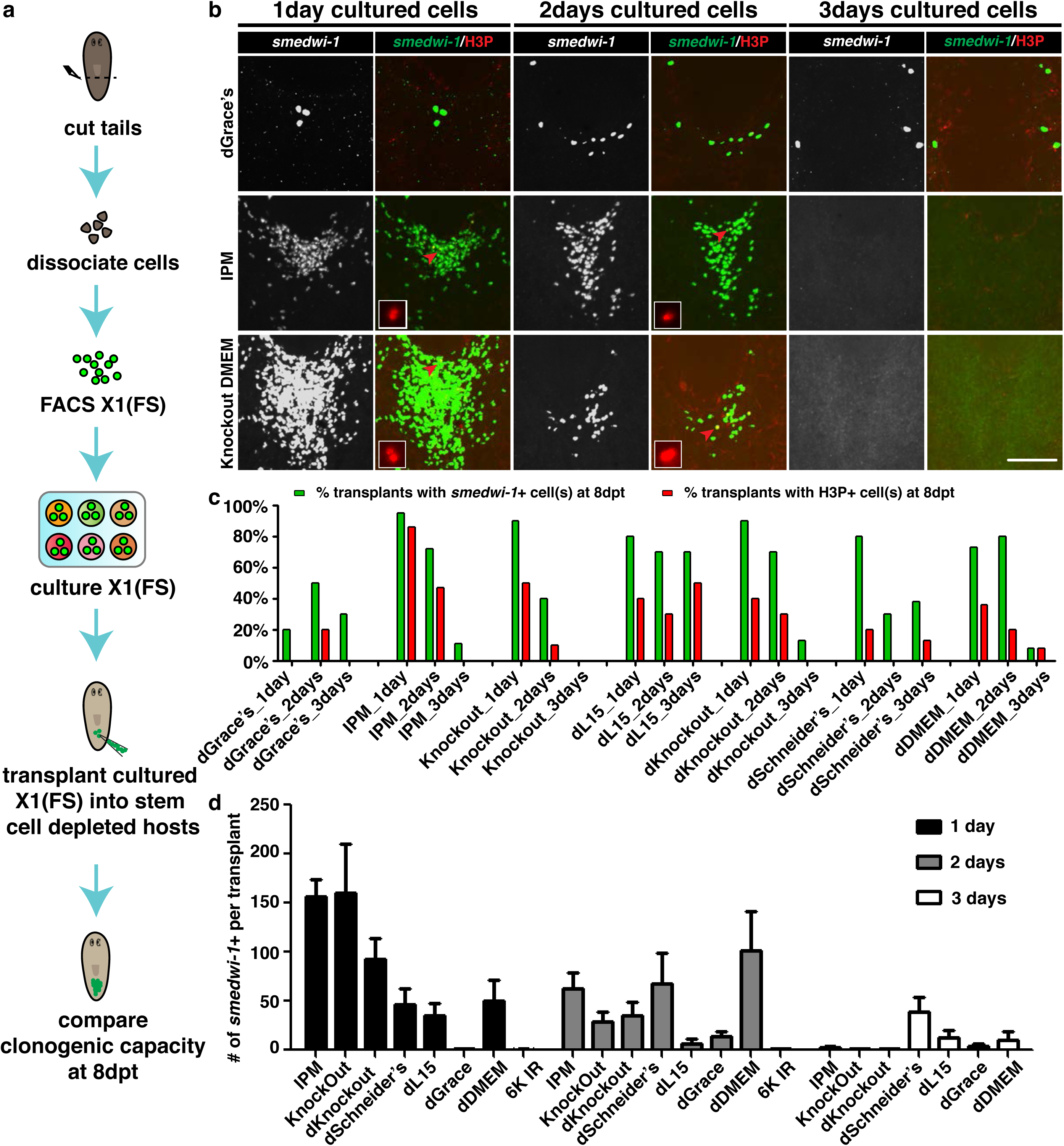
Cultured X1(FS) neoblasts expand after transplantation. **(a)** Flowchart showing steps of X1(FS) cell transplantation following *in vitro* culture. **(b)** Representative images showing colonies of *smedwi-1+* neoblasts at 8 days post-transplantation (dpt) cultured in the indicated conditions + 5% CO_2_. Only X1(FS) cells cultured in dGrace’s medium + 5% CO_2_ did not efficiently form colonies *in vivo*. Scale bar, 200 µm. **(c)** Percentage of hosts receiving X1(FS) cells cultured in indicated media + 5% CO_2_ for 1, 2, or 3 days that possessed *smedwi-1+* colonies (green bars) or H3P+ colonies (red bars) at 8 dpt. **(d)** Number of *smedwi-1+* neoblasts in colonies formed by X1(FS) cells at 8 dpt following culturing in indicated media + 5% CO_2_ for 1, 2, or 3 days. Ten to twelve animals assayed per condition.

### Cultured neoblasts can rescue stem cell-depleted planarian hosts

To evaluate the functional pluripotency of neoblasts cultured in these six media (IPM, KnockOut DMEM, dKnockOut DMEM, dL15, dSchneider’s, and dDMEM), rescue was assessed following bulk-cell transplantation. Genotyping PCR and restriction fragment length polymorphism (RFLP) assays were performed to test whether sexual hosts had been transformed into the asexual biotype following transplantation of the asexual neoblasts (Supplementary Fig. 5a) ^8^. For non-cultured, freshly collected X1(FS) cells, 30–50% of the lethally irradiated (6,000 rads) sexual *S. mediterranea* hosts were rescued (Fig. 3b, c, and Supplementary Fig. 5b, e). Next, X1(FS) cells cultured in the indicated media for 1, 2, or 3 days were transplanted into lethally irradiated hosts using the same method. X1(FS) cells cultured in IPM, dL15, and KnockOut DMEM for 1 and 2 days were capable of rescuing hosts devoid of stem cells (Fig. 3c and Supplementary Fig. 5c-e), of which X1(FS) cells cultured in KnockOut DMEM displayed the highest and most robust rescue efficiency (Fig. 3c and Supplementary Fig. 5e). Consistent with the clonogenic assay results, none of the X1(FS) neoblasts cultured for 3 days rescued lethally irradiated hosts. These data indicate that of all conditions tested, KnockOut DMEM +5% CO_2_ is the best one for maintaining pluripotent neoblasts in culture for up to 2 days. IPM and dL15 medium were also capable of maintaining pluripotent neoblasts in culture for up to 2 days albeit with reduced rates of irradiate animal rescue after transplantation (Fig. 3c and Supplementary Fig. 5e).

**Figure 3.**
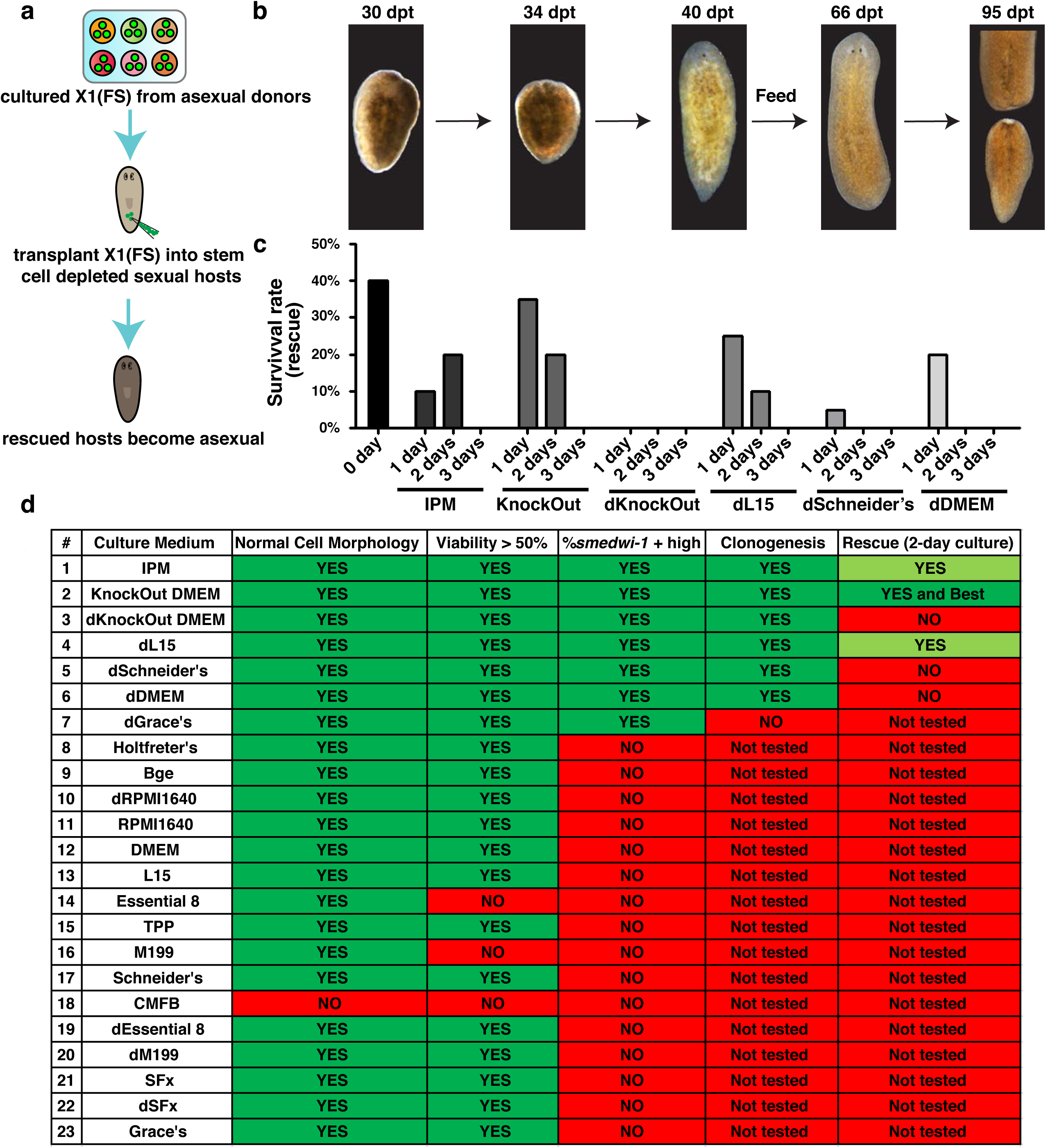
Cultured X1(FS) cells rescue neoblast-depleted planarians. **(a)** Flowchart illustrates steps of rescue assay using cultured X1(FS) cells. **(b)** Representative images showing rescue of lethally irradiated hosts following transplantation of freshly isolated X1(FS) cells, culminating in fission at 95 dpt. Scale bar, 200 µm. **(c)** Rescue rates for lethally irradiated hosts following the transplantation of X1(FS) cells cultured in the indicated media + 5% CO_2_ for 1, 2, or 3 days. Histogram indicates averaged percent from replicate experiments. Ten to twelve animals assayed per condition in each replicate experiment. **(d)** Summary of 23 cell culture media screen using the following criteria: cell morphology, cell viability, %*smedwi-1+* neoblasts, ability of transplanted cells to form colonies and expand *in vivo* (clonogenesis), and ability to rescue lethally irradiated animals (pluripotency). Overall, KnockOut DMEM was the most effective medium for maintaining pluripotent neoblasts in culture for 2 days.

In summary, we found that after screening 23 media followed by assaying 5 criteria (*i.e.*, viability, *smedwi-1* expression, cell division, clonogenic capacity and rescue efficiency of irradiated animals), three types of media (KnockOut DMEM, IPM, and dL15) were capable of maintaining pluripotent neoblasts *in vitro*. Of these three different media, KnockOut DMEM produced cultured neoblasts with the strongest performance across the multiple measured criteria, with IPM and dL15 medium performing slightly less well (Fig. 3d).

### Electroporation delivers fluorescent dextran into neoblasts

Following the successful optimization of *in vitro* culture conditions for the maintenance of pluripotent neoblasts, we next attempted to test conditions for the delivery of exogenous molecules into neoblasts, the next step required for developing transgenic methods for planarians. We first used dextran-FITC as a fluorescent indicator to screen suitable electroporation conditions for neoblasts labeled by Hoechst 33342 staining (Fig. 4a). We tested 52 electroporation programs and 10 different buffers using X1 cells ^25, 26^, and found that dextran-FITC was delivered into neoblasts most efficiently in IPM buffer with electroporation at 100-120V (Supplementary Table 2 and Fig. 4b-d). When similarly applying the electroporation method to X1(FS) cells, rather than Hoescht 33342 sorted X1 cells, dextran-FITC+ populations could only be detected with electroporation values of 110V and 120V. However, less than 6% of dextran-FITC+ X1(FS) cells were *smedwi-1+* neoblasts and virtually no *smedwi-1+* cells could be detected after 1 day culture in KnockOut DMEM +5% CO_2_ (Fig. 4e). Consistent with the drastic reduction in *smedwi-1+* cell viability post-electroporation, none of the donor X1(FS) cell populations subjected to more than 100V formed colonies following transplantation into lethally irradiated donors (Fig. 4f). We reasoned that the failure was likely due to the low purity of *smedwi-1+* neoblasts in X1(FS), which was even further reduced after electroporation. Therefore, it was necessary to develop a new strategy for neoblast isolation that would result in both higher clonogenic and pluripotent *smedwi-1+* cell enrichment than the X1(FS) sorting protocol.

**Figure 4.**
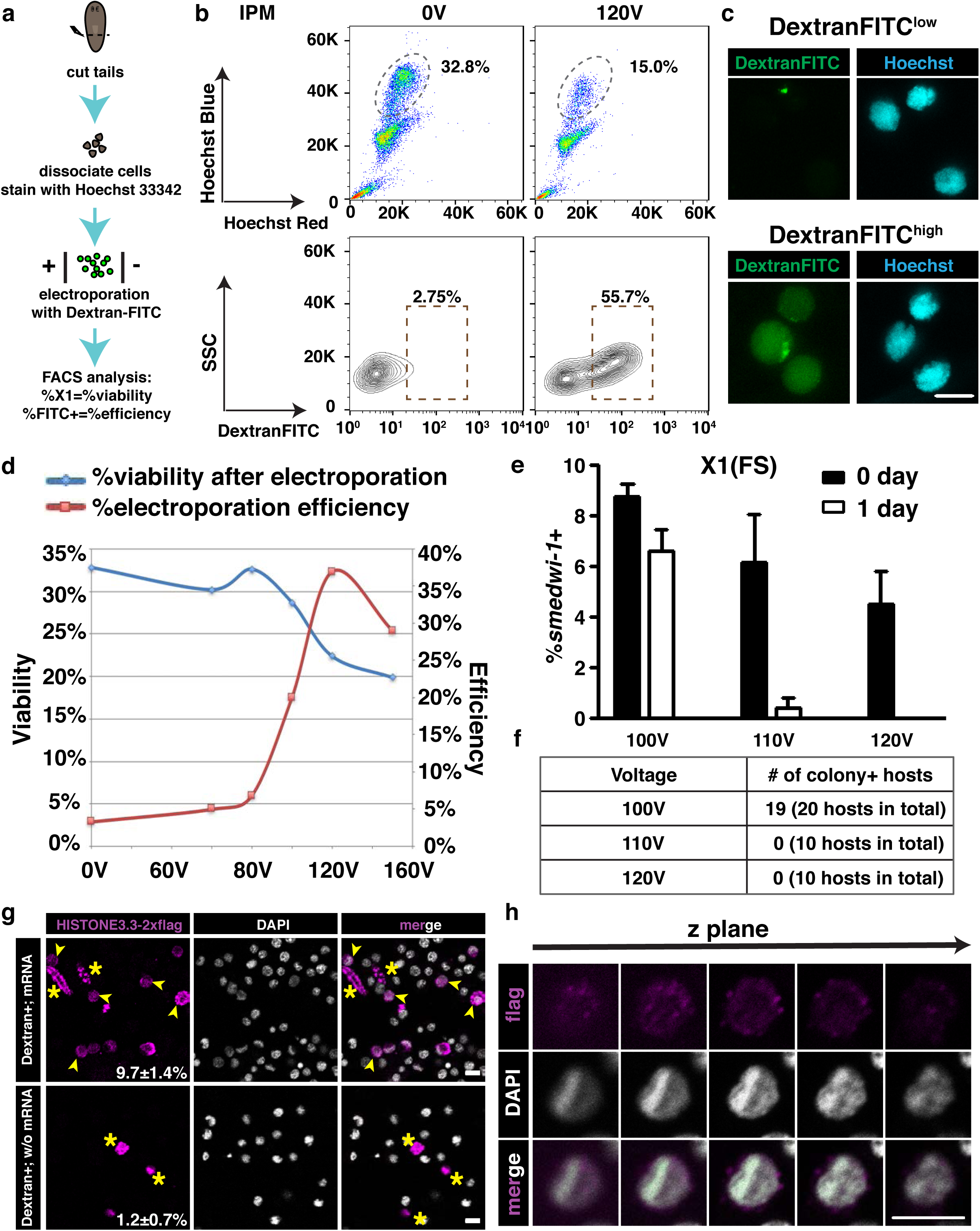
Electroporation can deliver Dextran-FITC into neoblasts. **(a)** Flowchart describing electroporation assay steps to screen for best conditions for cell viability and Dextran-FITC deilivery efficiency. **(b)** Plots of X1 viability (upper) and electroporation efficiency (lower) by using IPM as the electroporation buffer to deliver Dextran-FITC at 120V compared to 0 V controls. **(c)** Representative images of sorted Dextran-FITC^low^ and Dextran_FITC^high^ cells indicate successful delivery of Dextran-FITC at 120V. **(d)** Viability (blue) and electroporation efficiency (red) on X1 cells after electroporation using IPM as electroporation buffer. **(e)** %*smedwi-1*+ neoblasts in X1(FS) cells after 100V, 110V, and 120V electroporation immediately (black column) and after 1 day of culture in KnockOut DMEM + 5% CO_2_ (white column). Four random fields assayed per condition. p<0.05 for 120 V. N>40. **(f)** Electroporated X1(FS) cells receiving greater than 100 V failed to form colonies following transplantation. Ten animals assayed per condition. **(g)** Representative images of electroporated X1(FS) with (upper panel) or without (lower panel) *Smed-histone3.3-2×flag* mRNA. Arrowheads: anti-FLAG+ nucleated cells. Stars: anti-FLAG+ enucleated cells. Scale bar, 20 µm. **(h)** Z-stack images of an nucleated anti-FLAG+ cell. Scale bar, 10 µm.

We also tested whether X1(FS) can express exogenously delivered mRNA in current culture conditions. We cloned a planarian endogenous gene, *Smed-histone3.3* and fused with two copies of flag tag (2×flag). After electroporation and one day of culture, cells electroporated with *Smed-histone3.3-2×flag* mRNA had more anti-FLAG staining positive cells (9.7 ± 1.4%) than electroporated cells without mRNA (1.2±0.7%) (Fig. 4g). Even though the anti-FLAG antibody stained enucleated cells, nuclear localization signal in nucleated cells suggested successful expression of *Smed-histone3.3-2×flag* mRNA (Fig. 4g and 4h). Even though we did not detect the signal of Smed-HISTONE3.3-2×FLAG in *smedwi-1*+ cells (Supplementary Fig. 6), these data encouraged us to further enrich for neoblasts to optimize cell culture conditions.

### A new flow cytometry protocol using SiR-DNA and Cell Tracker improves yield of clonogenic, pluripotent, transplantable neoblasts

To enrich for neoblasts, we tested three major types of cell-permeable DNA stains to enrich cycling neoblasts at G2/M cell cycle phases (DRAQ5, Vybrant DyeCycle, and SiR-DNA). DRAQ5 staining remained cytotoxic similarly to Hoechst 33342. Vybrant DyeCycle staining failed to unambiguously discriminate among distinct neoblast cell cycle phases by flow cytometry. However, the recently developed DNA stain, SiR-DNA ^29^ proved to have low toxicity and enriched *smedwi-1*+ neoblasts to a ratio ∼60% (Fig. 5a, b, f and Supplementary Fig. 7). Comparison of *smedwi-1*+ and *smedwi-1*- cell morphology in the isolated populations showed that *smedwi-1*- cells were generally smaller than *smedwi-1*+ cells (Fig. 5b). To discriminate between small and large cells in the SiR-DNA+ population, the cytoplasmic dyes Cell Tracker Green (CT) and Calcein AM (CAM) were tested in combination with SiR-DNA in neoblast isolation (Fig. 5c, d). Using a dual dye staining strategy resulted in a significant increase in neoblast enrichment, as judged by *smedwi-1*+ ISH (Fig. 5e, f); SiR-DNA/Cell Tracker Green costaining performed comparably to Hoechst 33342 staining for enriching *smedwi-1*+ neoblasts (Fig. 5e, f). We termed this new sorted cell population SirNeoblasts. Unlike neoblasts derived from Hoechst 33342 sorts, SirNeoblasts proliferated *in vitro* and underwent colony expansion *in vivo* after transplantation into lethally irradiated planarians (Fig. 5g). Facilitated by SiR-DNA staining of DNA, the separation dynamics of chromosomes in dividing SirNeoblasts were observed *in vitro* (Supplementary Movies 3-5), confirming the occurrence of *bona fide* cell division in the tested culture condition. Importantly, no noticeable difference in colony sizes was observed at 7 dpt between X1(FS), single (SiR-DNA), and double dye (SiR-DNA/CT) stained populations (Fig. 5g). Finally, both freshly isolated SirNeoblasts and those cultured in KnockOut DMEM +5% CO_2_ for one day were capable of rescuing lethally irradiated planarians at frequencies comparable to those seen with X1(FS) cells (Fig. 3c and Fig. 5h). Based on these results, we conclude that SiR-DNA/CT dual labeling-based cell sorting can be used to isolate clonogenic, pluripotent neoblasts that can be maintained in primary culture and serve as donor cells in transplantation assays. To further characterize the SirNeoblasts, we stained SirNeoblasts with Hoechst 33342 to analyze their cell cycle. However, co-staining of SiR-DNA and Hoeschst 33342 resulted in a failure to detect SiR-DNA staining. We then tested whether Hoeschst 33342 can stain SiR-DNA stained cells, and found that Hoechst 33342 can replace the staining of SiR-DNA, which showed the cell cycle distribution of SirNeoblasts consisted of ∼17.89% at G1, 13.02% at S, and ∼69.09% at G2/M cell cycle phases (Supplementary Fig. 7). This reversible staining of SiR-DNA may also explain the reason why SirNeoblasts can proliferate after staining unlike Hoechst 33342 stained X1 cells.

**Figure 5.**
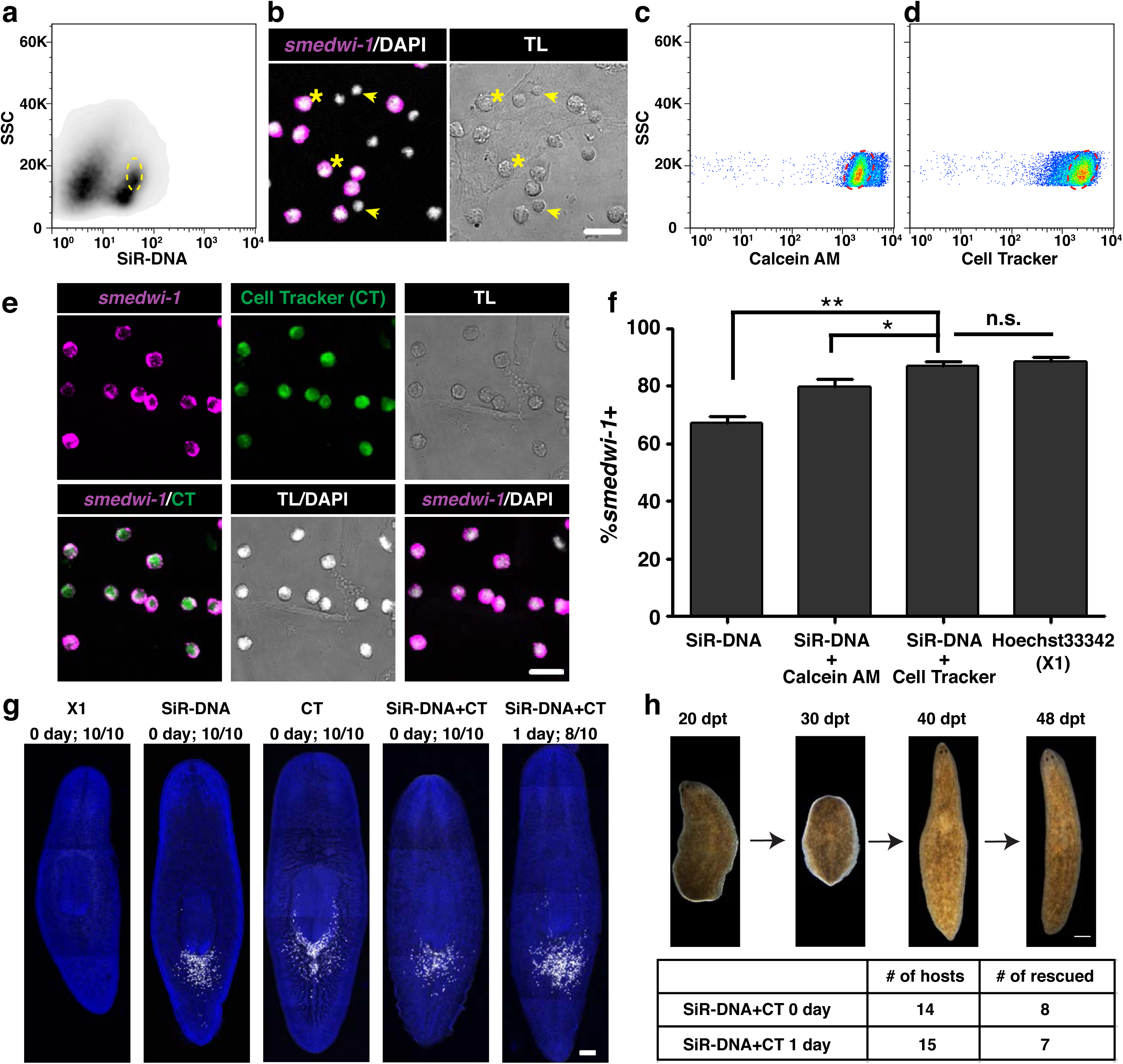
SiR-DNA plus Cell Tracker staining and cell sorting protocol enriches for clonogenic, pluripotent *smedwi-1+* neoblasts. **(a)** Plots showing the gate used to isolate SiR-DNA+ cells for *smedwi-1* ISH. **(b)** *smedwi-1* ISH on isolated cells from the SiR-DNA+ gate shown in Supplementary Fig. 7a. *smedwi-1*- cells (arrows) were generally smaller than *smedwi-1*+ cells (stars). Scale bar, 20 µm. **(c-d)** Plots showing the gates used to isolate SiR-DNA+, calcein-AM+ cells **(c)** and SiR-DNA+, Cell Tracker Green+ cells **(d)** for *smedwi-1* ISH. **(e)** *smedwi-1* ISH for SIR-DNA+ neoblasts populations indicated in **(c)**. Scale bar, 20 µm. **(f)** %*smedwi-1*+ neoblasts in indicated FACS isolated populations. SiR-DNA and Cell Tracker Green dual staining enriches for *smedwi-1+* neoblasts (SirNeoblasts) comparably to the Hoechst 33342 stained X1 population. *, p<0.05. **, p<0.01. n.s., no significance. Four random fields assayed per condition. N>70. **(g)** Representative images showing the clonogenic capacity of transplanted neoblasts obtained using different FACS isolation protocols. No noticeable difference in the colony expansion was observed among single and double dye staining populations at 7dpt. Scale bar, 200 µm. Ten animals assayed per condition. **(h)** Rescue efficiency of fresh and 1-day cultured SirNeoblasts. CT: cell tracker green.

Next, we determined conditions to optimize electroporation efficiency and viability for SirNeoblasts (Fig. 6a). Consistent with previous studies, electroporation at 110V-120V was required for dextan-TMR entry into SirNeoblasts (Fig. 6b, c). As expected, *smedwi-1*+ cells were more abundant in the 110 V and 120V electroporated SirNeoblasts compared to X1(FS) cells, and some electroporated SirNeoblasts persisted for one day in culture (Fig. 6d). Importantly, 110V – 120V electroporated SirNeoblasts were capable of forming colonies and rescuing lethally irradiated hosts upon transplantation (Fig. 6e, f). However, 120V electroporations resulted in comparably fewer irradiated hosts being rescued after SirNeoblast transplantations (Fig. 6e, f), indicating that high voltages may have a negative impact on pluripotency.

**Figure 6.**
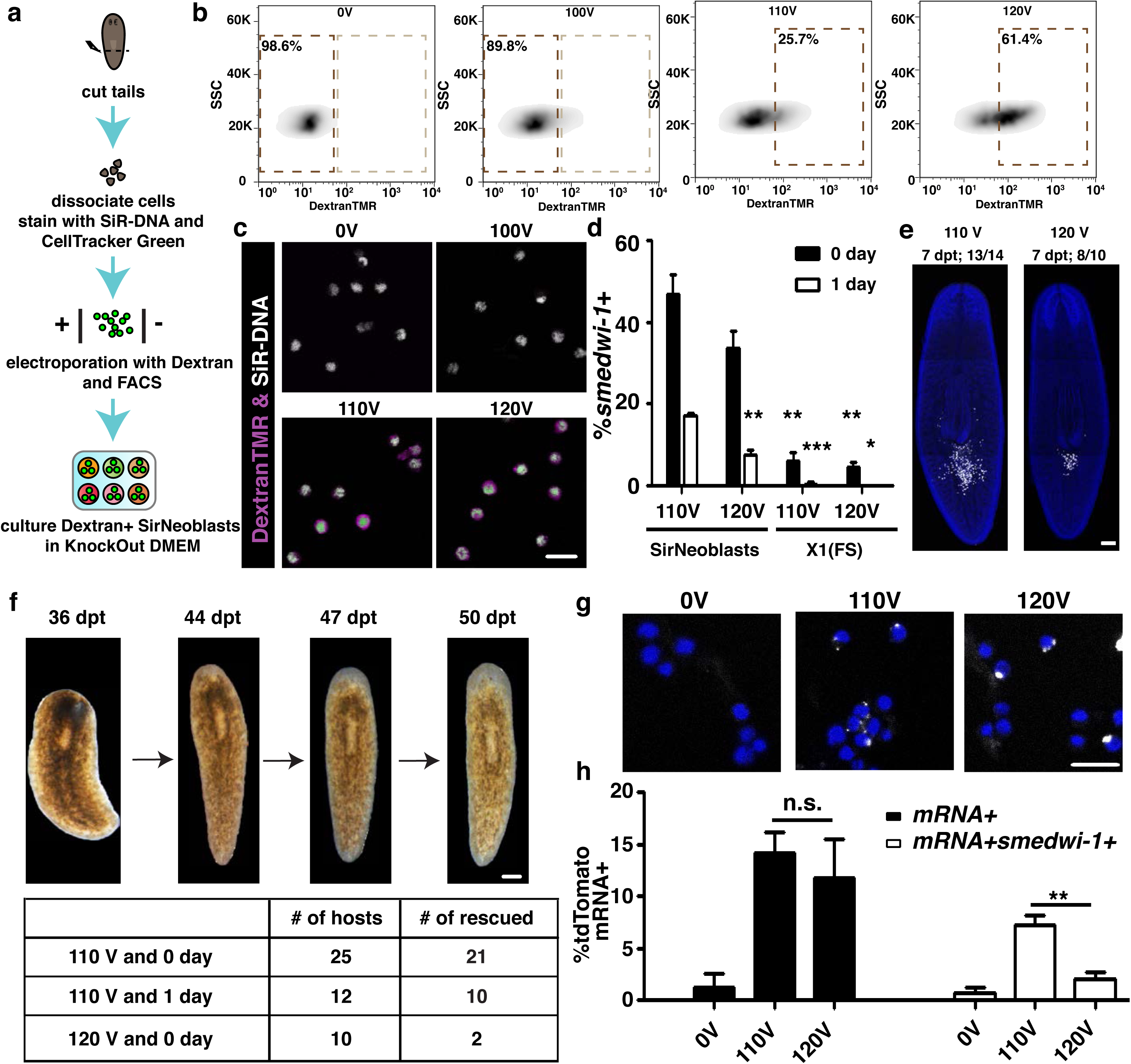
SirNeoblasts can be used for exogenous mRNA electroporation. **(a)** Flowchart presenting the steps of neoblast electroporation using SirNeoblasts. **(b)** Plots showing electroporation efficiency of SirNeoblasts at 100V, 110V and 120V compared to 0V. **(c)** Neoblasts after electroporation of Dextran-FITC showing 100% isolation of positive cells after electroporation at 110V and 120V. All SirNeoblasts were free of Dextran-FITC without electroporation treatment. Scale bar, 20 µm. **(d)** Percentage of *smedwi-1*+ cells after electroporation, suggesting a relative high ratio of neoblasts after electrporation by using SirNeoblasts compared to X1(FS) in Fig. 4e. Four random fields assayed per condition. *, p< 0.05 (120V SirNeoblasts vs. 120V X1(FS) at 1 day). **, p<0.005 (110V SirNeoblasts vs. 120V SirNeoblasts at 1 day, 110V SirNeoblasts vs. 110V X1(FS) at 0 day, and 120V SirNeoblasts vs. 120V X1(FS) at 0 day). ***, p<0.001 (110V SirNeoblasts vs. 110V X1(FS) at 1 day). **(e)** Representative images showing the colony expansion of electroporated SirNeoblasts after transplanation Scale bar, 200 µm. N=14 for 110V and =10 for 120 V. **(f)** Rescue efficiency of electroporated SirNeoblasts. Scale bar, 200 µm. **(g)** Representative images showing the mRNA signals (white dots) in cells 1 day after 110V and 120V electroporation. Scale bar, 20 µm. **(h)** Percentage of total cells and *smedwi-1+* cells containing mRNA 1 day after 110V and 120V electroporation. n.s.: not significant. ** < 0.01.s

### Exogenous mRNA delivered by electroporation can be successfully expressed in SirNeoblasts

To assess whether exogenous mRNA could be delivered into SirNeoblasts using the described electroporation conditions, *tdTomato* mRNA was added to the electroporation reaction along with Dextran. Dextran positive SirNeoblasts were sorted and cultured in KnockOut DMEM + 5% CO_2_. To determine whether mRNA was successfully delivered, we probed cells via FISH 20 hours after electroporation. *tdTomato* mRNA signal was detected in both 110V and 120V electroporated cells, suggesting a successful delivery of exogenous mRNA into SirNeoblasts (Fig. 6g, h). However, costaining with *smedwi-1+* revealed that not all *tdTomato* mRNA+ cells retained neoblast identity in culture. The number of sorted SirNeoblasts responded similarly to X1 and X1(FS) cells with respect to electroporation in that the number of cells positive for both tdTomato mRNA and *smedwi-1* expression was significantly higher after 110V electroporation than after 120V (Fig. 6h). This result indicates that under the conditions tested, 110V electroporation may be the most suitable to both introduce exogenous, charged molecules such as RNA into neoblasts, while maintaining their viability and potency.

Unfortunately, expression of tdTomato was not detected by either microscopy or antibody staining. Two possibilities were suspected: 1) The culure condition is not good enough to support the translation of the delivered mRNA; 2) There is an unknown mechanism that prevents the translation of the delivered mRNA. A recent discovery in *C. elegans* indicated that endogenous piRNAs can target on the exogenous transgene sequences and prevent their translation ^30^. Similarly, planarian neoblasts contain abundant PIWI and piRNAs. We thus hypothesize that a similar piRNA targeting mechanism may exist in planarian neoblassts, which may prevent the translation of the delivered mRNAs. We tested this hypothesis by synthesizing multiple mRNAs encoding the fluorescent protein mCherry in which conservative nucleotide substitutions were introduced in order to minimize potential pairing of the exogenous mRNA with endogenous piRNAs, as was recently described in *C. elegans* ^30^. The synthetic mCherry mRNAs were tested via electroporation into SirNeoblasts (Fig. 7a). Significantly, we found one mCherry mRNA construct that resulted in robust mCherry+ cultured SirNeoblasts (Fig. 7b-e, twice with high expression, five times with medium/low expression, ten times without expression). Even though we have yet to fully comprehend the mechanisms that may be underpinning piRNA targeting in neoblasts, the successful expression results indicated that the culture and electroporation conditions defined in our study are capable of maintaining neoblasts in culture capable of retaining both pluripotency (Figures 5g, h and 6e, f) and translational activity (Figure 7b). Although the efficiency by electroporation is low for mRNA delivery, our current study is focused on developing a reliable method for culturing neoblasts. Increasing the efficiency of delivery for mRNA and testing Cas9 and guide RNAs is clearly necessary and will require further studies.

**Figure 7.**
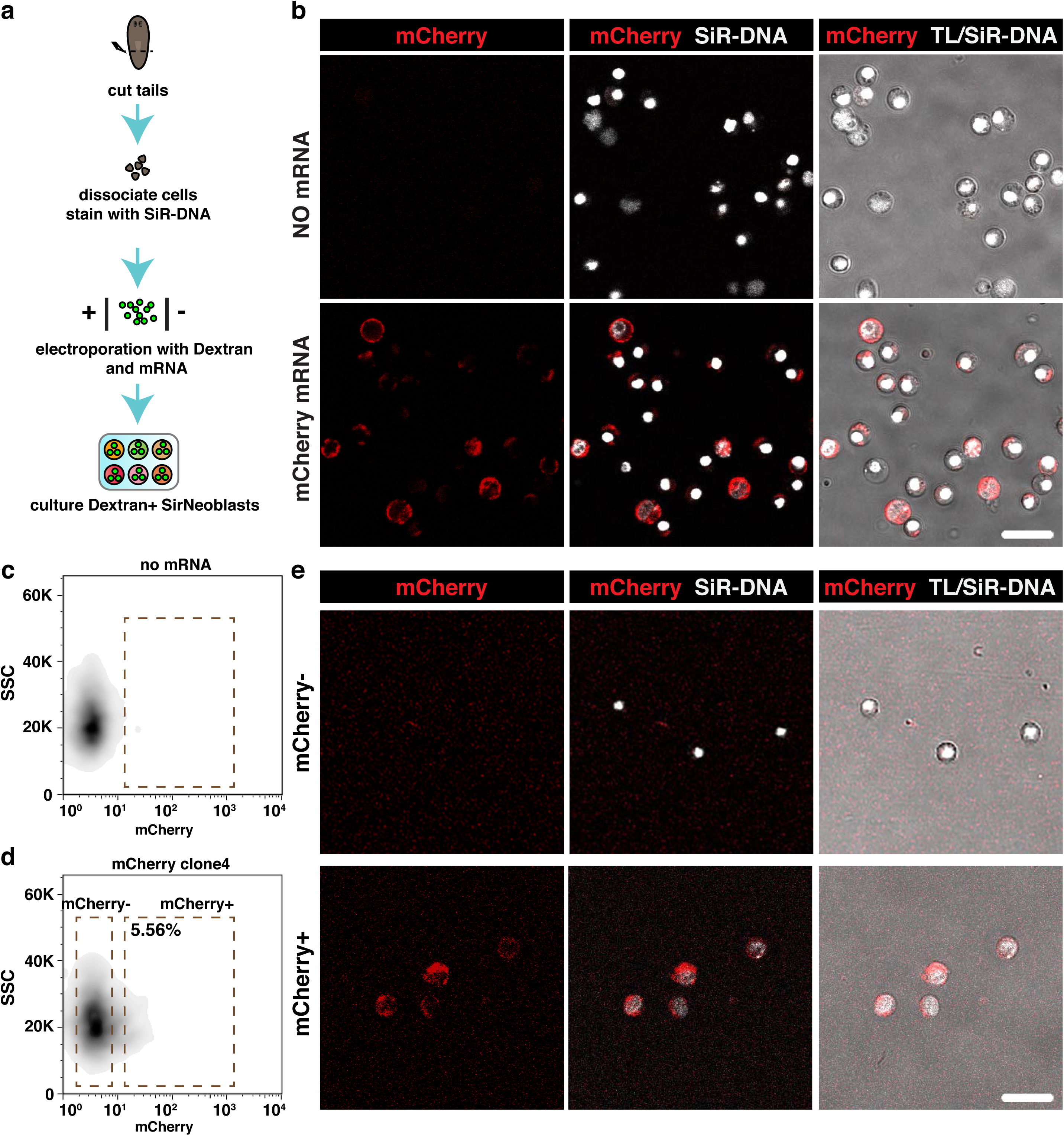
mCherry mRNA is expressed in SirNeoblasts. **(a)** A flowchart describes steps of SirNeoblast electroporation using mCherry mRNA. **(b)** Representative images of mCherry mRNA electroporated SirNeoblasts cultured in KnockOut DMEM for 1 day. Upper: electroporated SirNeoblast without mRNA in culture. Lower: mCherry mRNA electroporated SirNeoblasts in culture. Scale bar, 20 µm. **(c)** Plot showing no mCherry expression after 110V electroporation without mCherry mRNA. **(d)** Plot showing ∼5% mCherry+ cells after 110V electroporation with mCherry mRNA. **(e)** Representative images of cells from mCherry- population in (upper row) and mCherry+ population in (lower row). Cells from mCherry+ population showed obvious mCherry localization in cytoplasm. Scale bar, 20 µm.

In summary, we defined a novel FACS isolation strategy and primary cell culture conditions capable of maintaining clonogenic, pluripotent neoblasts *in vitro* that are compatible with transplantation, repopulation and rescue of lethally irradiated hosts. In addition, we optimized electroporation conditions that successfully introduced fluorescent dextran and exogenous mRNA into clonogenic, pluripotent neoblasts. These technical milestones are prerequisites for the successful generation of transgenic planarians.

## Discussion

Past efforts to culture planarian cells have been unable to convincingly demonstrate that pluripotent neoblasts could be maintained in culture ^23, 24, 31, 32^. Here, we provide definitive molecular and functional evidence that pluripotent neoblasts can be maintained *in vitro*. This technical advance facilitated the first real-time observation of neoblast cell division within the first 24 hours after cell culture *in vitro* (Supplementary Movies 1 and 2) and the first demonstration that exogenous molecules, including fluorescent conjugated dextrans and mRNA, can be delivered into planarian cells. This method establishes the required foundation for future transgenic and genome editing technique development in planarians, and opens exciting new avenues for a systematic investigation of the biology of a naturally occurring population of pluripotent adult stem cells.

### The vital fluorescent dye SiR-DNA improves purification of pluripotent neoblasts

Prior to this study, The use of Hoechst 33342 staining has been broadly adopted for isolating cycling neoblasts (X1 cells) by FACS. However, X1 cells labeled with this nuclear dye cannot proliferate *in vivo* following their transplantation into irradiated hosts. We sought to overcome this limitation by testing alternative DNA dyes, such as DRAQ5 and Vybrant Dye Cycle, yet they resulted in cytotoxicity and failed to unambiguously discriminate between distinct neoblast cell cycle phases by flow cytometry. However, we found that unlike other vital DNA dyes tested, the recently developed SiR-DNA dye ^29^ was not cytotoxic, and when combined with Cell Tracker Green staining, significantly improved pluripotent neoblast yields by flow cytometry. Together with cell subtype-specific antibodies, SiR-DNA may allow for more specific dissection of the pluripotent neoblast population by facilitating the isolation and functional characterization of different neoblast subpopulations. Furthermore, this reagent may prove useful for the isolation of viable proliferating cells in other organisms.

### Neoblast cell culture paves the way for transgenesis in planarians

Transgenesis in planarians has been lacking for decades. Without a planarian-specific positive control, it has been difficult to determine why exogenous nucleic acids fail to be translated when introduced into neoblasts. Isolated neoblasts provide an obvious proving ground for delivery of exogenous materials. While neoblast transplantation can be performed immediately after delivering exogenous molecules, the uncertain viability of neoblasts during and after transplantation made this strategy ineffective. The cell culture system we have developed makes it possible to trace and study each cell following delivery of exogenous materials *in vitro*. First, it allows for ease of screening of constructs using a small number of cells under conditions where neoblast purity and viability are well-established. Second, when introducing transformed cells into lethally irradiated hosts to monitor behavior *in vivo*, we can enrich for positive cells via FACS prior to transplantation, minimizing the effects of cell-cell competition in a heterogeneous donor cell population. Hence, neoblasts cultured using the methods described here lend themselves accessible for testing a diversity of delivery methods. For instance, custom-engineered liposomes were shown to facilitate the transfection of double-stranded RNA and anti-miRNAs into planarian cells *in vivo* ^33^. As such, it should be possible to use liposomes to deliver larger molecules and genome-editing tools in an effort to obtain higher neoblast transfection efficiency and further improve the likelihood of producing transgenic animals. Thus, our methodology not only stands to facilitate cell transformation, but may also play a key role in efforts to establish long-term culture systems and/or cell lines.

### piRNA silencing mechanism for transgenes may be of broad occurrence across metazoans

Efforts to generate transgenic planarians span several decades with little to no success reported thus far. The reasons for this state of affairs have been generally associated with technical limitations of both culture conditions and delivery methods of exogenous nucleic acids into neoblasts. Little consideration, however, has been given to the possibility that such prolonged failure may be underscored by unknown aspects of neoblast biology. Given that neoblasts are the *de facto* units of selection in planarians and that the viability of these animals heavily depends on the proper function and health of neoblasts, strong positive selection for evolving robust mechanisms to protect the genome of these cells should be expected.

piRNAs are small non-coding RNAs that have been shown to be essential for safeguarding genome integrity by silencing transposable elements ^34^. However, it is also known that many piRNAs do not map to transposable element sequences in various animals, including mice, *C. elegans* and planarians ^35–37^. In fact, the function of these piRNAs remains largely unknown. Additionally, it has also been known for decades that transgenes with foreign sequences can be frequently silenced in the germline of *C. elegans* ^38^. Recent studies have begun to shed light on piRNA function in both *Drosophila* ^39^ and *C. elegans* ^40^. It was recently reported that the repression of transgenes in the germline of fruitflies could be lifted by using a UAS-promoter free of interference by Hsp70 piRNAs as silenced ^39^. Also, it is known that the PIWI protein PRG-1 is required for the silencing phenomenon observed in the germline of *C. elegans*, suggesting a function of piRNAs in this process ^40^. More recently, it was discovered that a mechanism targeting transgene sequences introduced into the syncytial ovary of *C. elegans* involves piRNAs, and that a sequence-based strategy to bypass transgene silencing by these small non-coding RNAs allowed expression of exogenously added genes in the germline of this animal ^30^.

Given the ancestral origin of PIWI proteins and piRNAs, we hypothesized that similar mechanisms may be operating in planarian neoblasts. Our current study showed that exogenous mRNAs in which predicted piRNA targeting sequences were changed overcame siencing and allowed the translation of the reporter protein (Figure 7b). However, we do not yet fully understand the piRNA recognition rules in *S. mediterranea*. The size of planarian piRNAs are ∼32nt, in contrast to ∼22 nt in *C. elegans* ^37, 41^, so the models prediciting targeting of piRNA in nematodes do not fully transpose to planarians. Additionally, planarians have at least three PIWI proteins ^25, 37^, raising another question as to which PIWI proteins may or may not be required for producing piRNAs that may potentially target exogenous nucleic acids. Definitive experiments to test this hypothesis are necessary and future and ongoing research will help in resolving these issues and testing and refining our predictive piRNA targeting models in the hopes of producing the most stable exogenous nucleic acid molecules for introduction into neoblasts.

### A method for mechanistic studies of neoblast proliferation and differentiation *in vitro*

The paucity of cell culture conditions for invertebrates in general, and planarians in particular, has hampered our ability to test and identify factors directly regulating the functions of neoblasts, a remarkably abundant and pluripotent adult stem cell population. For example, our understanding of how extracellular growth factors modulate neoblast proliferation is still in its infancy. In planarians, several of these factors have been shown to have important functions in neoblast proliferation or homeostasis. For instance, knockdown of *smed-neuregulin (nrg)-7* or *smed-insulin-like peptide-1* impairs neoblast proliferation *in vivo* ^13, 42^. We hypothesize that addition of these factors, or potentially other purified extracellular growth factors, may boost neoblast proliferation *in vitro*. However, no *in vitro* culture system had been developed to test this hypothesis. With the methods and results presented here open the door to test the effects of planarian extracellular extracts or purified extracellular growth factors from planarian species on the proliferation and maintenance of neoblasts. Additionally, our protocols lend themselves to initiate a systematic comparison of the metabolomics of cultured neoblasts with those found *in vivo*. Such studies will aid in further optimization of culture conditions and may ultimately lead to the controlled manipulation of cell metabolism to predictably regulate neoblast proliferation and differentiation *in vitro*.

### Defining the neoblast niche in planarians

The existence of a niche that supports the proliferation and differentiation of neoblasts has been previously proposed ^43^. This hypothesis has been supported by indirect evidence ^13, 44, 45^. However, the molecular and cellular nature of the niche is largely unknown. Transplant experiments carried out in this study showed that a limited number of neoblasts can be maintained in the transplanted location and may continue their proliferation and differentiation. Because of the limited number of cells surviving after transplantation, dissecting the cellular microenvironment of transplanted neoblasts is likely to be a promising context for a mechanisitic characterization of the proposed neoblast niche. Together with sublethal irradiation assays, the cell culture tools reported here should afford us the opportunity to understand how pluripotency and cell fate may be cell- and non-cell autonomously regulated in a highly regenerative context.

### A framework for establishing cell culture in new research organisms

Since the development of cell lines in the 1950s ^46^, cell culture has enabled scientists to study fundamental aspects of cell biology. In recent years, the number of research organisms being employed to address and discover new biology has steadily increased. However, a comparatively small number of cell types have been successfully cultured *in vitro*, particularly for invertebrates. The current study systematically screened the majority of published cell culture media and optimized culture conditions for planarian neoblasts. Thus, the systematic development of cell culture methods reported here not only advances the study of cell bbiology in the highly regenerative planarian *S. mediterranea*, but should also facilitate the establishment of culture methods for other species, particularly underrepresented invertebrate research organisms.

## Experimental Procedures

### Planarian care and irradiation treatment

Asexual (Clone CIW4) and sexual (Clone S2F1L3F2) strains of *Schmidtea mediterranea* were maintained in Montjuïc water at 20°C as previously described ^8, 20^. Animals were starved for 7–14 days before each experiment. Animals exposed to 6,000 rads of γ rays were used as transplant hosts ^8^. After transplantation, hosts were maintained in Montjuïc water with 50 µg/ml Gentamicin (GEMINI, 400-100P). For transplant rescue experiments, host animals were maintained in 3.5 cm Petri dishes (1 worm/dish), and Montjuïc water was changed every 2–3 days.

### Cell collection and culture

X1(FS) cells were collected as previously described with minor modifications ^8, 25^. Tails from planarians (>8 mm in length) were macerated in Calcium Magnesium free buffer with 1% Bovine Serum Albumin (CMFB) (see Recipe in Table S1) for 20–30 min with vigorous pipetting every 3–5 min. After maceration, dissociated cells were centrifuged at 290g for 10min. Cells were then resuspended in IPM with 10% Fetal Bovine Serum for either Hoechst 33342 or SiR-DNA staining. To gate the X1(FS) cells, the X1 population from a control sample stained with 0.4 mg/ml Hoechst 33342 (ThermoFisher Technologies, H3570) was used to define the forward scatter/side scatter gate. To obtain SirNeoblasts, dissociated cells were stained with SiR-DNA (1µM, Cytoskeleton Inc., CY-SC007) and CellTracker Green CMFDA Dye (2.5µg/ml, Thermo Fisher Technologies, C7025) for 1 hour and 10 min sequentially. Cells were sorted with an Influx sorter using a 100-µm tip. For time-lapse imaging experiments, X1(FS) cells were incubated in either 5 mL of the indicated culture medium per well in 6-cm dishes (MatTek, P35G-1.5-14-C) or in 1 mL of the indicated culture medium per well in a 24-well plate (MatTek, P24G-1.5-13-F). For other experiments, X1(FS) were incubated in 50 µL of the indicated culture medium per well in 384-well plates (Greiner bio-one, 781090). Cells were cultured in indicated media containing 5% Fetal Bovine Serum (Sigma-Aldrich, F4135) at 22°C, +/− 5% CO_2_. Dishes and multi-well plates were pre-coated with poly-D-lysine (50µg/ml, BD Biosciences).

### *In situ* hybridization and antibody staining

Whole-mount *in situ* hybridization was carried out as previously described ^13, 47–49^. For ISH on cultured cells, cell culture plates were centrifuged in an Eppendorf horizontal centrifuge (Centrifuge 5810 R) at 300 *g* × 3 min. Cells were fixed with 3.7% formaldehyde (Sigma-Aldrich, F8775) or 4% paraformaldehyde (Electron Microscopy Sciences, 15710) for 20 min. After washing with 1× PBS, cells were hybridized with riboprobes at 56°C for at least 15 h. After washing with 2× SSC and 0.2× SSC, cells were incubated with anti-digoxigenin-POD (Roche Diagnostics, 11207733910) or anti-fluorescein-POD (Roche Diagnostics, 11426346910) at room temperature for 2 h. After washing with 1× PBS/0.3% TritonX-100, the signal was developed using tyramide-conjugated Cy3 (Sigma-Aldrich, PA13101) or Cy5 (Sigma-Aldrich, PA15101).

Anti-phospho-Histone H3 (Ser10) (H3P) antibody (1:1,000, Abcam, ab32107) and Alexa 555-conjugated goat anti-rabbit secondary antibodies (1:1,000, Abcam, ab150086) were used to stain proliferating cells at the G2/M phase of the cell cycle.

### Annexin V staining

Fifty microliters of cultured cells were re-suspended and stained with 2.5 µl of Annexin V FITC Conjugate (BioLegend, 640905) at room temperature for 15 min. After washing twice with IPM + 10%FBS, cells were subjected to *smedwi-1* ISH. Thereafter, anti-fluorescein-POD (Roche Diagnostics, 11426346910) was used to stain Annexin V for apoptotic and dead cells detection.

### Cell transplantation

X1(FS) cells collected by flow cytometry were transplanted into irradiated hosts (6,000 rads) as previously described with minor modifications ^8^. Approximately 1 µL of an X1(FS) cell suspension (5,000 cells/µL) was injected into either the post-pharyngeal midline (of asexual CIW4 hosts) or the post-gonopore midline (of sexual S2F1L3F2 hosts) at 0.75–1.0 psi (Eppendorf FemtoJet) using a borasilicate glass microcapillary (Sutter Instrument Co., B100-75-15).

### mRNA synthesis and electroporation

mRNAs used for electroporation were prepared following the standard protocols in the mMESSAGE mMACHINE T7 ultra Transcription Kit (ThermoFisher Technologies, AM1345) and the Ambion RNA Purification Kit (ThermoFisher Technologies, AM1908). tdTomato mRNA was transcribed from the linearized plasmid pcDNA3.1(+)-tdTomato. mCherry and T7 promoter sequences were cloned into the pIDT vector and synthesized by IDT Inc. Primers used to amplify the template were 5′-CAGATTAATACGACTCACTATAGG-3′ and 5′-ACTGATAATTAACCCTCACTAAAG-3′.

To screen electroporation conditions, cells from four tail fragments were suspended in 20 µL electroporation buffers following Heochast 33342 staining. 20 µg Dextran-FITC (ThermoFisher Technologies, D3306) were mixed with cells and loaded into a 1mm electroporation cuvette for BTX ECM830 electroporator or a 12-well electroporation strip for Lonza 4D electroporator. The buffer SE, SG, SF, P1-5 were electroporation buffers in Lonza Cell Line and Primary Cell 4D-Nucleofector Optimization kits (V4XC-9064 and V4XP-9096). Cell viability and electroporation efficiency were assessed using an Influx sorter.

For exogenous mRNA electroporation, ∼1×10^8^ cells were suspended in 50 µL IPM following SiR-DNA staining. 50 µg Dextran-FITC and ∼5 µg mRNA were mixed with cells and loaded into a 1mm electroporation cuvette. BTX ECM830 electroporator was used to apply a 110 V and 1 millisecond square wave pulse to deliver dextran-FITC and mRNA into planarian cells. Dextran-FITC+ SirNeoblasts were purified using an Influx sorter and cultured in KnockOut DMEM + 5%FBS.

### Microscopy and time-lapse imaging

The Celigo imaging cell cytometer (Celigo, Inc.) and the Falcon 700 confocal microscope were used to take pictures of X1(FS) and SirNeoblasts following ISH. Celigo or ImageJ software was used for quantitative analyses. Time-lapse imaging of cultured cells was performed using a Nikon Eclipse TE2000-E equipped with Perfect Focus and a Plan Fluor ELWD 20X/0.45 NA Ph1 objective. Micro-manager was used to control the microscope and Hamamatsu Orca R2 CCD ^50^. Multiple positions were acquired at 5-min intervals for 24–48 h. *In situ* hybridization samples were imaged with a Nikon Eclipse Ti equipped with a Yokogawa W1 spinning disk head and a Prior PLW20 Well Plate loader. Several slides were prepared at once and then loaded and processed automatically using a combination of Nikon Elements Jobs for all robot and microscope control and Fiji for object-finding and segmentation. Slides were imaged at low magnification and objects identified before re-imaging tiled z-stacks using a Plan Apo 10X 0.5NA air objective. Tiled images were stitched, projected, and *smedwi-1+* puncta were counted using custom macros and plugins in Fiji.

### Generation of optimized mCherry sequence

mCherry candidate sequences were generated by means of a custom python script. Amino acid sequences were back translated to 21 nucleotide sequences from 7 amino acid words at a time. Each potential nucleotide sequence was screened against a list of known piRNAs to generate the sequence with the fewest piRNA matches. A piRNA match consists of no more than a single G/T mismatch in the 6 nucleotide seed region (positions 2-7 of a piRNA) ^30^. Additional G/T mismatches were scored as .5 and other mismatches as 1. Only the first 21 basepairs of the piRNAs were aligned. The highest scoring piRNA determined the score for that potential nucleotide sequence. The 21 nucleotide sequence with the lowest score was retained. The script was run with four alternate coding tables. The “all” coding table contained all possible codons for each amino acid. The “smed” coding table contained only those codons known to be most used in *S. mediterraea* ^51^. “lowgc” contained only those codons with the fewest G or C nucleotides. “highgc” contained only those codons with the most G or C nucleotides. The “highgc” sequence is shown in Figure 7. The other three sequences as well as 5 additional sequences generated by shuffling the four generated sequences and one sequence generated by backtranslating the amino acid sequence with sms failed to show fluorescence ^52^.

## Data availability

All codes used for plugins in Fiji are available at:https://github.com/jouyun. All original data underlying this manuscript can be accessed from the Stowers Original Data Repository at: http://www.stowers.org/research/publications/libpb-1281. All reagents are available from the corresponding author upon reasonable request.

## Statistical analyses

Microsoft Excel and Prism 6 were used for statistical analysis. Mean ± s.e.m. is shown in all graphs. Unpaired two-tailed Student’s *t*-test was used to determine the significant differences between two conditions. p < 0.05 was considered a significant difference.

## Acknowledgments

We thank I. Wang and P. Reddien for assistance with the transplantation technique. We thank all members of Sánchez Lab, especially J. Jenkin and C. Guerrero for animal maintenance and irradiation assistance, L. C. Cheng and E. Duncan for technical help, and B. Benham-Pyle, E. Davies, and S. Elliot for comments on the manuscript. We acknowledge all members of the Reptile & Aquatics, Molecular Biology, Cytometry, and Microscopy Core Facilities at the Stowers Institute for Medical Research for technical support. This work was supported by NIH R37GM057260 to A.S.A. A.S.A. is a Howard Hughes Medical Institute and a Stowers Institute for Medical Research Investigator.

## Author contributions

K.L. and A.S.A. conceived the project, designed experiments, analyzed data, and wrote the manuscript. K.L. performed all experiments and data acquisition. S.A.M. performed the time-lapse imaging experiments and analyzed raw spinning-disk imaging data. E.J.R. and H.-C.L. designed the variant sequences for mCherry.

## Figure Legends

**Supplementary Figure 1.**
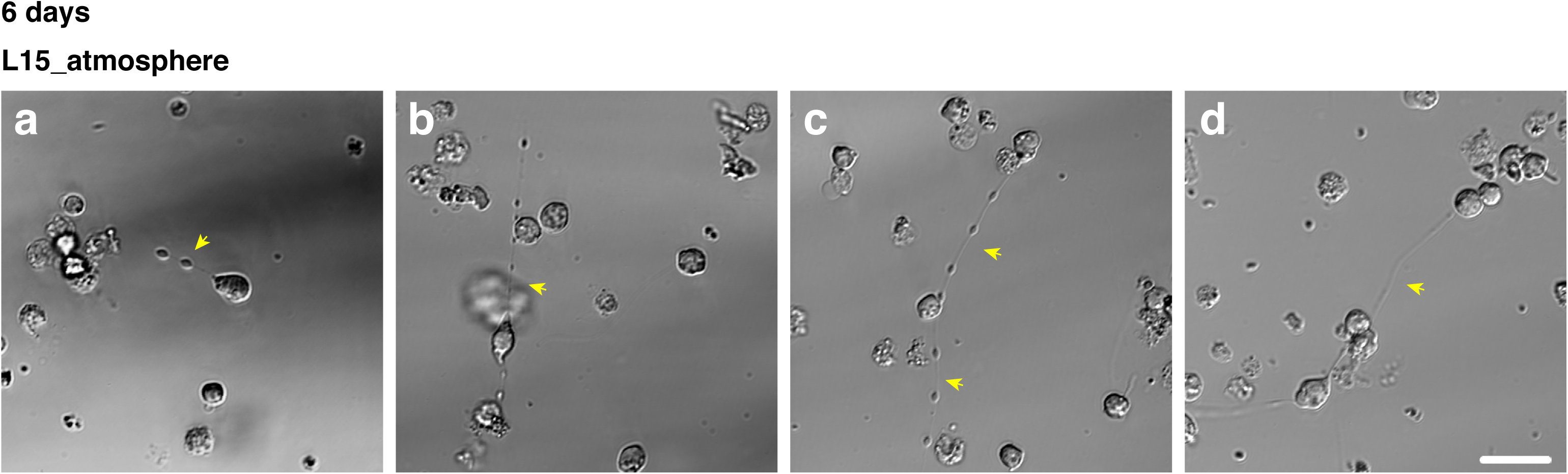
X1(FS) cells cultured in L15 extend long cellular processes. **(a–d)** Four representative images showing long cellular processes from cells after 6 days of culture in L15 without 5% CO_2_. Scale bar, 20 µm.

**Supplementary Figure 2.**
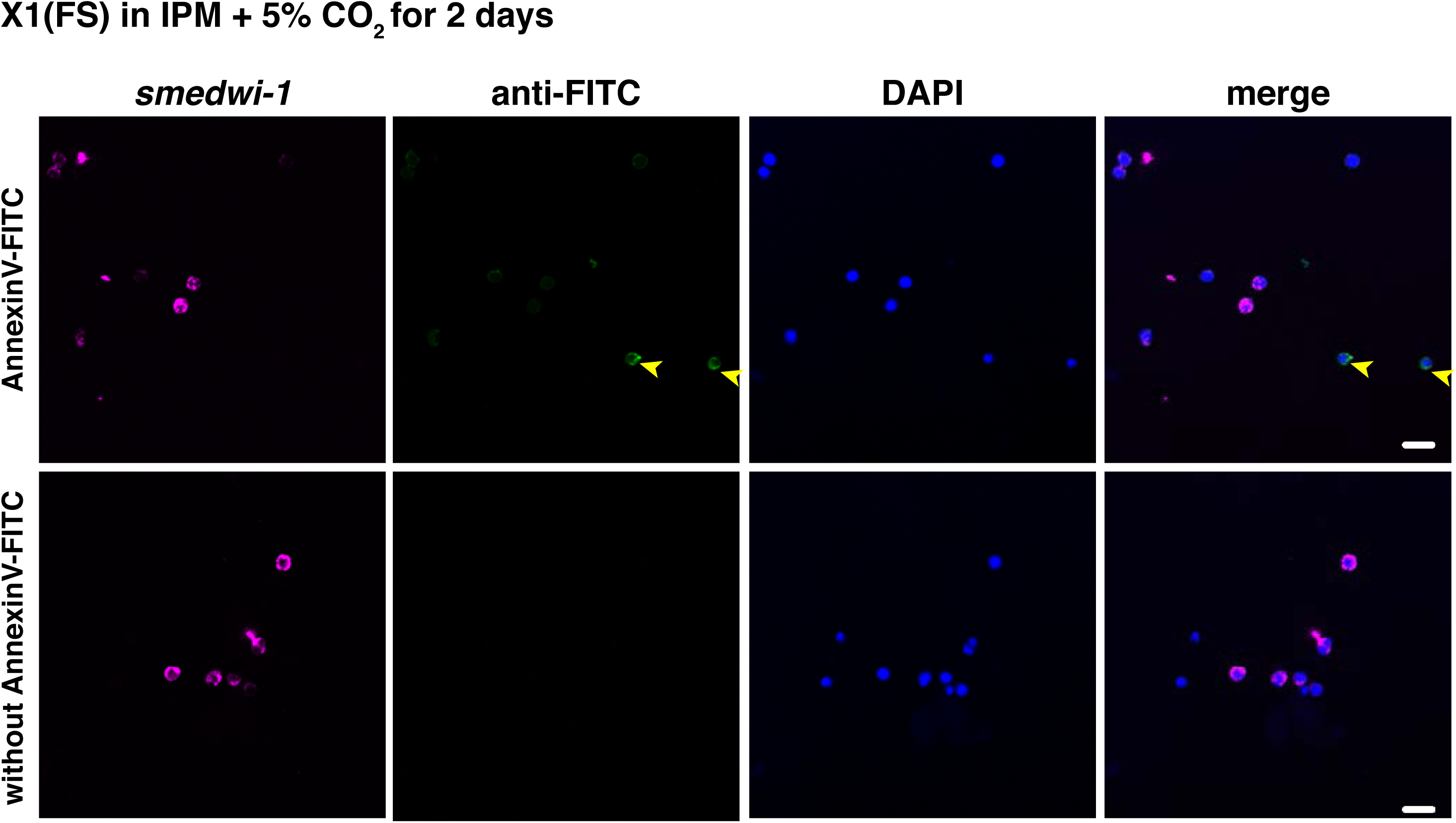
s*medwi-1*+ X1(FS) neoblasts are viable. X1(FS) cells were cultured in IPM + 5% CO_2_ for 2 days. Representative images of apoptotic cells (Annexin V, green, arrowheads) co-stained with the pan-neoblast marker *smedwi-1* (magenta), n=37. Two independent replicate experiments were performed. No co-labeling was observed, suggesting neoblasts examined in study were viable. Scale bar, 20 µm.

**Supplementary Figure 3.**
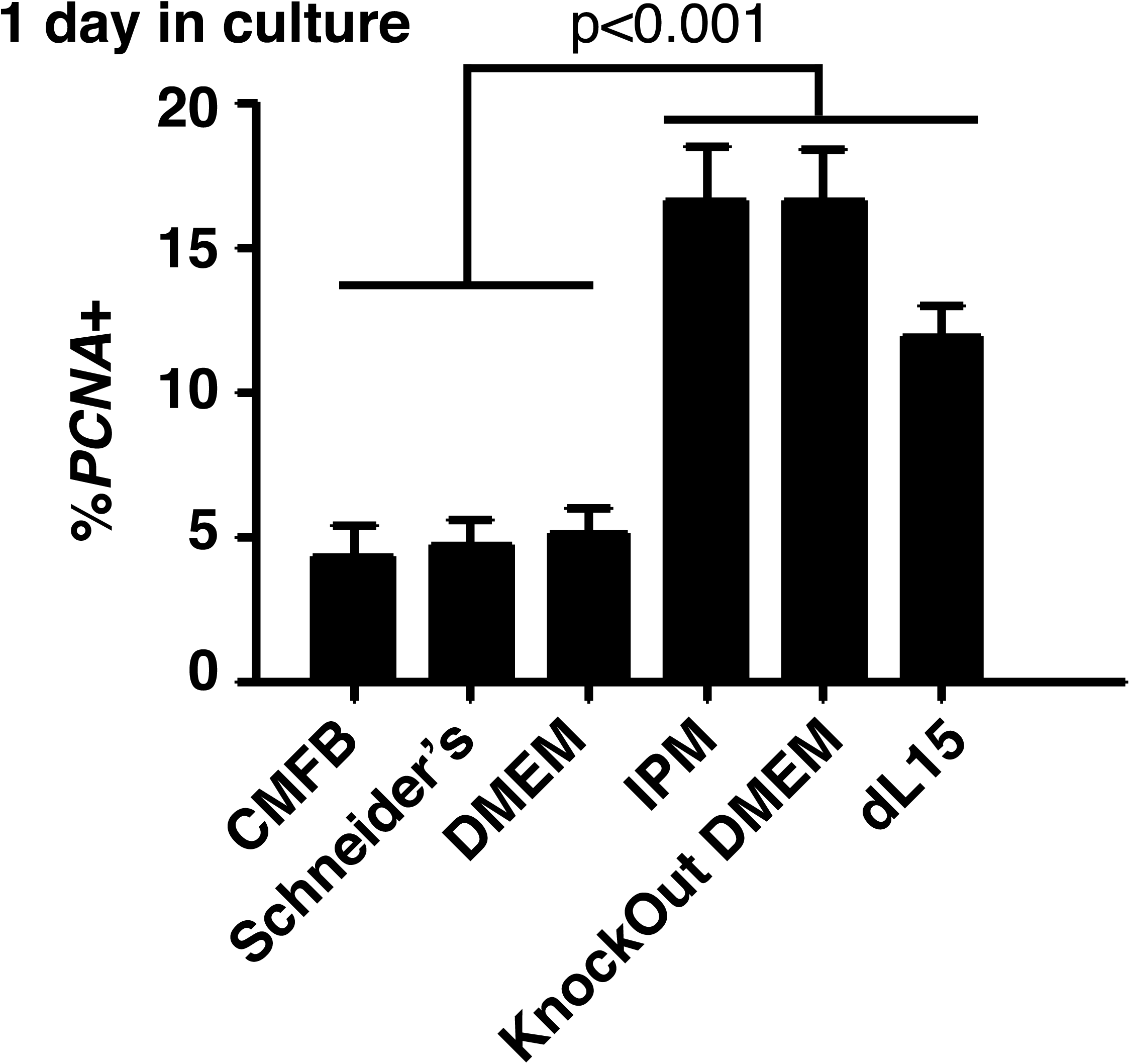
IPM, Knockout DMEM, and dL15 maintain more *PCNA*+ cells. Percentage of *smedwi-1+* neoblasts after 1 day of culture in indicated media + 5% CO_2_.

**Supplementary Figure 4.**
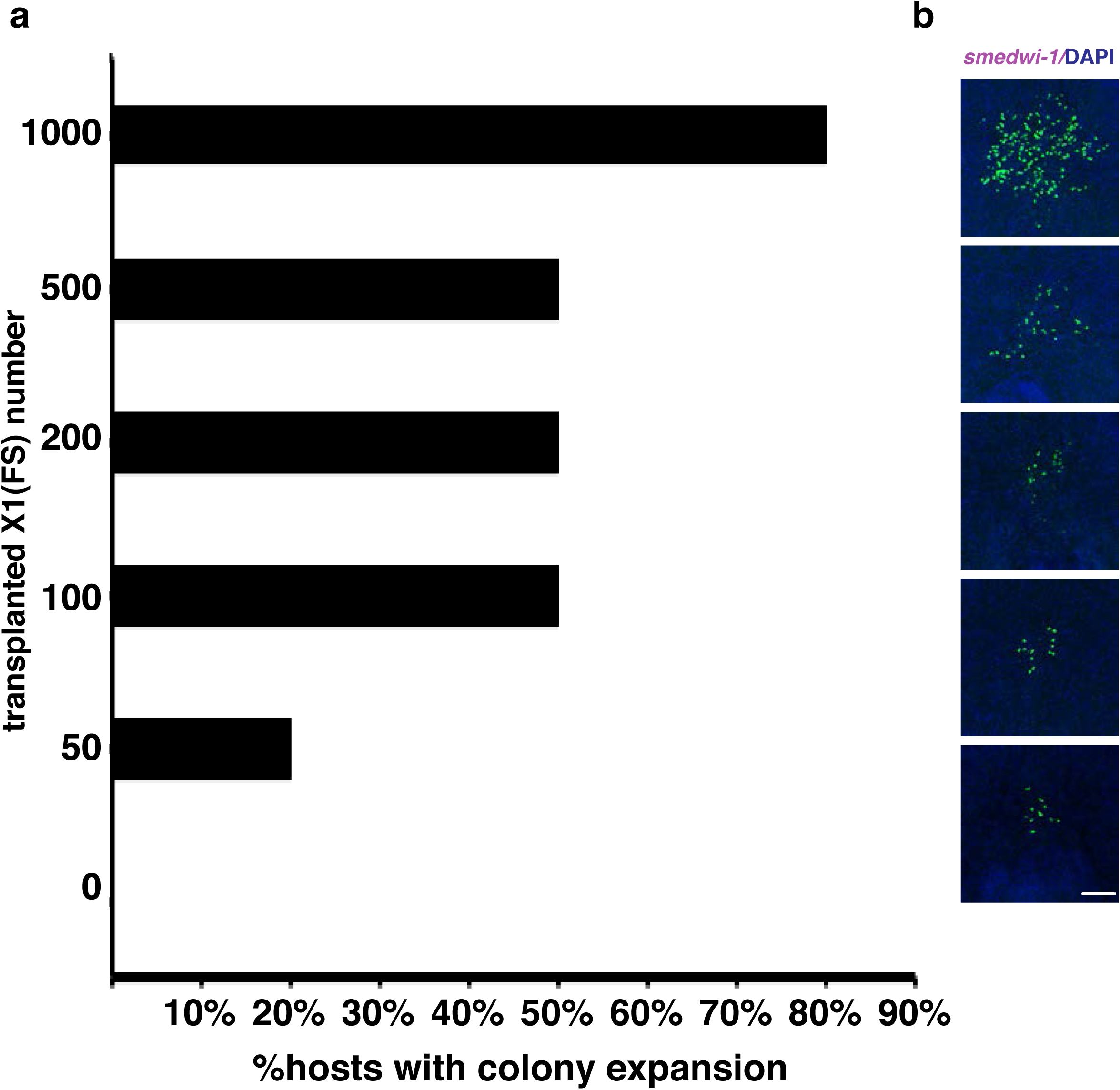
Determining the number of X1(FS) cells needed for efficient colony expansion. **(a)** Percentage of lethally irradiated hosts displaying robust neoblast colony expansion following transplantation with the indicated numbers of sorted X1(FS) cells. At 7 days post-transplantation (dpt), > 80% of all hosts displayed colony expansion when 1,000 X1(FS) were transplanted. **(b)** Representative images of hosts transplanted with X1(FS) cells at 7 dpt. *smedwi-1*+ neoblasts: green. DAPI: blue. Scale bar, 200 µm. Ten animals assayed per condition.

**Supplementary Figure 5.**
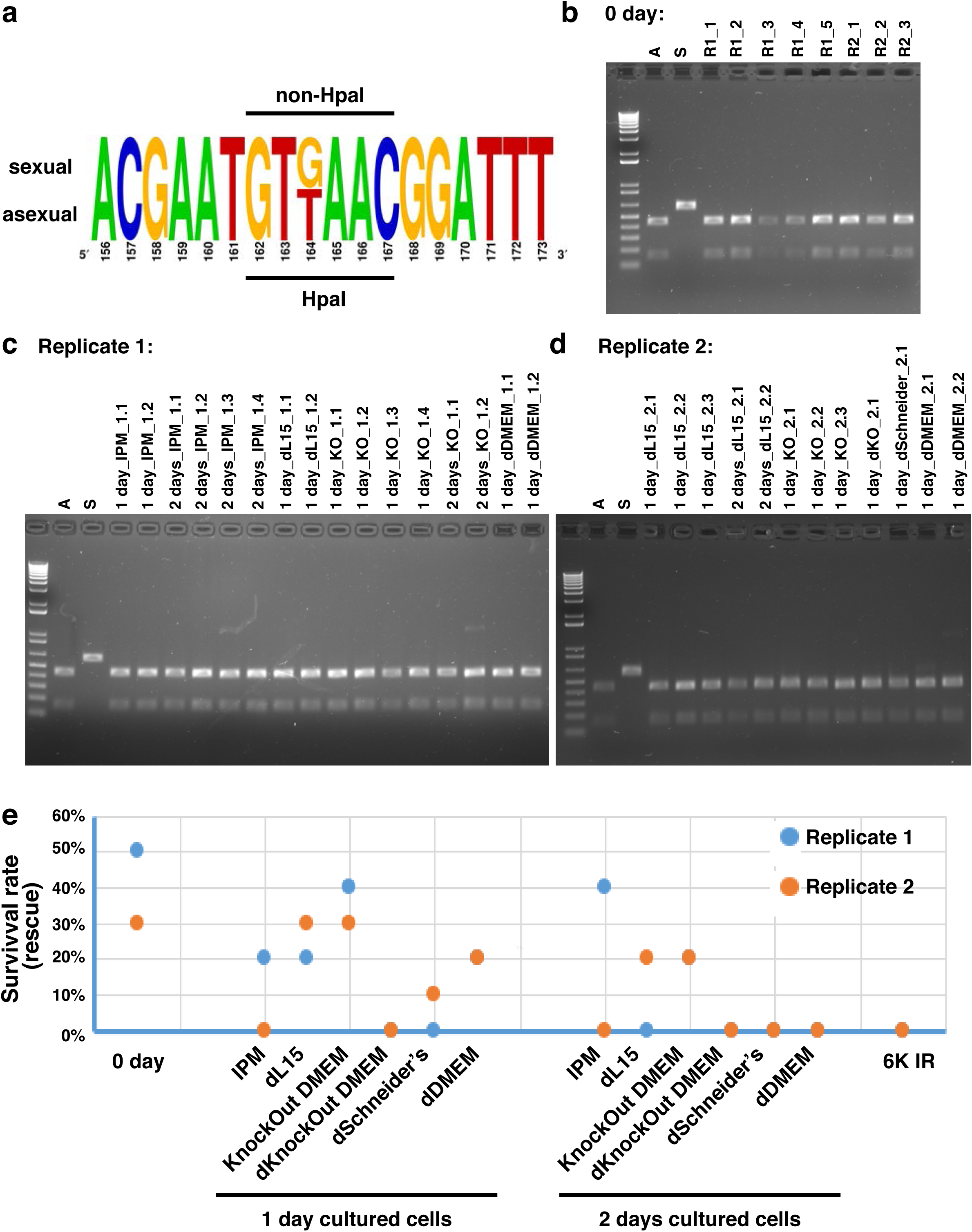
Sexual hosts are rescued and reconstituted by transplantation of cultured asexual X1(FS) cells. **(a)** Sequence showing the HpaI enzyme restriction site, which was used to distinguish between the asexual (donor) and sexual (host) biotypes by RFLP analyses. **(b)** RFLP data showing rescue of lethally irradiated sexual worms transplanted with freshly collected, non-cutured X1(FS) cells. **(c-d)** RFLP data showing rescue of lethally irradiated sexual worms transplanted with 1 and 2 day cultured X1(FS) cells. Data from two independent experiments shown replicate 1 (panel c); replicate 2 (panel d). **(e)** Rescue rates for lethally irradiated hosts following transplantation of X1(FS) cells cultured in the indicated media + 5% CO_2_ for 1 or 2 days. None of the conditions rescued lethally irradiated hosts after 3 days. Blue and orange dots show value of rescue rate from replicate experiments, respectively.

**Supplementary Figure 6.**
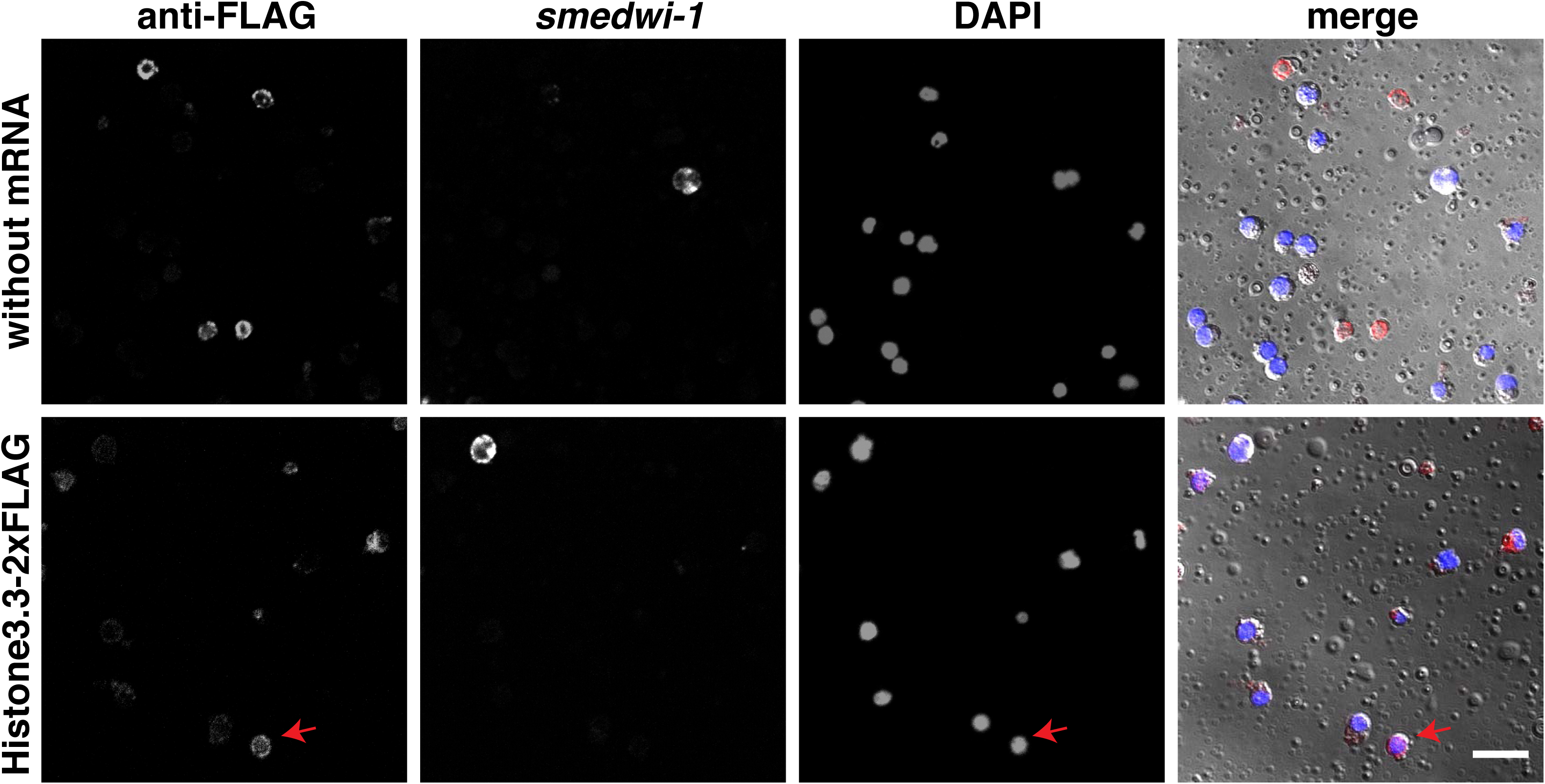
No expression of exogenously delivered *Smed-histone3.3-2*×*flag* mRNA in *smedwi-1*+ cells. Representative images of electroporated X1(FS) without (upper panel) or with (lower panel) *Smed-histone3.3-2×flag* mRNA. Cells cultured in Knockout DMEM + 5% CO_2_ for 1 day were stained with *smedwi-1* riboprobe and anti-FLAG antibody. Arrowhead: anti-FLAG+ nucleated cells. Scale bar, 20 µm.

**Supplementary Figure 7.**
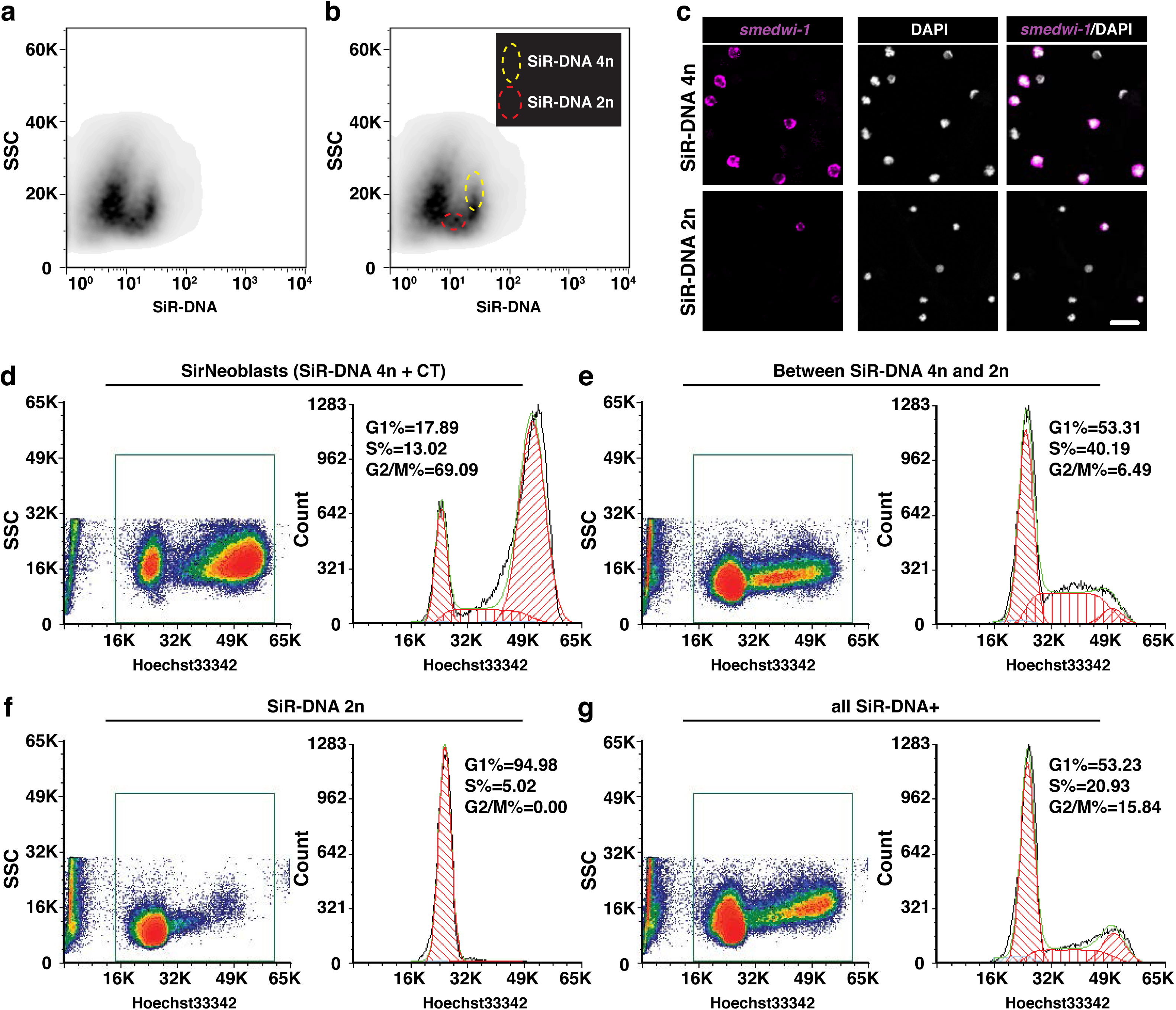
Compare SiR-DNA sorted cells. **(a)** A plot showing how SiR-DNA-stained cells are displayed without gates in the flow cytometry analysis using SiR-DNA versus side scater. **(b)** A plot showing how gates were defined to isolate two SiR-DNA staining cell populations based on DNA content (SiR-DNA 4n and 2n). **(c)** *smedwi-1* in situ staining for neoblasts in two isolated cell populations based on DNA content as indicated in (b). SiR-DNA 4n population contains 56.4%±2.6% *smedwi-1*+ neoblasts (also see Fig. 4f) compared to 26.8%±3.2% in SiR-DNA 2n population, p value = 0.0017. Scale bar, 20 µm. **(d-g)** Plots showing the cell cycle distribution of SirNeoblasts (SiR-DNA 4n + CT) (d), cells between SiR-DNA 4n and 2n (e), SiR-DNA 2n (f), and all SiR-DNA+ cells (g). Sorted cells were stained with Hoechst 33342. Hoechst 33342+ (square gate) cells were analyzed for cell cycle distribution.

